# Epigenetic signature and key transcriptional regulators of human antigen-specific type 1 regulatory T cells

**DOI:** 10.1101/2024.03.07.582969

**Authors:** Alma-Martina Cepika, Laura Amaya, Colin Waichler, Mansi Narula, Michelle M. Mantilla, Benjamin C. Thomas, Pauline P. Chen, Robert Arthur Freeborn, Mara Pavel-Dinu, Jason Nideffer, Matthew Porteus, Rosa Bacchetta, Fabian Müller, William J. Greenleaf, Howard Y. Chang, Maria Grazia Roncarolo

**Affiliations:** Division of Hematology, Oncology, Stem Cell Transplantation and Regenerative Medicine, Department of Pediatrics, Stanford University School of Medicine, Stanford, CA, USA; Institute for Stem Cell Biology and Regenerative Medicine, Stanford University School of Medicine, Stanford, CA, USA; Department of Dermatology, Stanford University School of Medicine, Stanford, CA, USA; Division of Infectious Diseases and Geographic Medicine, Stanford University School of Medicine, Stanford, CA, USA; Center for Definitive and Curative Medicine, Stanford University School of Medicine, Stanford, CA, USA; Integrative Cellular Biology and Bioinformatics, Saarland University, Saarbrücken, Germany; Department of Genetics, Stanford University School of Medicine, Stanford, CA, USA; Center for Personal Dynamic Regulome, Stanford University School of Medicine, Stanford, CA, USA; Howard Hughes Medical Institute, Stanford University, Stanford, CA 94305

## Abstract

Human adaptive immunity is orchestrated by effector and regulatory T (Treg) cells. Natural Tregs arise in the thymus where they are shaped to recognize self-antigens, while type 1 Tregs or Tr1 cells are induced from conventional peripheral CD4^+^ T cells in response to peripheral antigens, such as alloantigens and allergens. Tr1 cells have been developed as a potential therapy for inducing antigen-specific tolerance, because they can be rapidly differentiated *in vitro* in response to a target antigen. However, the epigenetic landscape and the identity of transcription factors (TFs) that regulate differentiation, phenotype, and functions of human antigen-specific Tr1 cells is largely unknown, hindering Tr1 research and broader clinical development. Here, we reveal the unique epigenetic signature of antigen-specific Tr1 cells, and TFs that regulate their differentiation, phenotype and function. We showed that *in vitro* induced antigen-specific Tr1 cells are distinct both clonally and transcriptionally from natural Tregs and other conventional CD4^+^ T cells on a single-cell level. An integrative analysis of Tr1 cell epigenome and transcriptome identified a TF signature unique to antigen-specific Tr1 cells, and predicted that IRF4, BATF, and MAF act as their transcriptional regulators. Using functional genomics, we showed that each of these TFs play a non-redundant role in regulating Tr1 cell differentiation, suppressive function, and expression of co-inhibitory and cytotoxic proteins. By using the Tr1-specific TF signature as a molecular fingerprint, we tracked Tr1 cells in peripheral blood of recipients of allogeneic hematopoietic stem cell transplantation treated with adoptive Tr1 cell therapy. Furthermore, the same signature identified Tr1 cells in resident CD4^+^ T cells in solid tumors. Altogether, these results reveal the epigenetic signature and the key transcriptional regulators of human Tr1 cells. These data will guide mechanistic studies of human Tr1 cell biology and the development and optimization of adoptive Tr1 cell therapies.

## Introduction

Regulatory T cells (Tregs) play a critical role in human health by inducing and maintaining tolerance to endogenous and exogenous antigens^1^. Type 1 regulatory T (Tr1) cells are Tregs that arise from effector CD4^+^ T cells (Teff) in response to repeated antigen stimulation in a tolerogenic environment, e.g., in the presence of immunoregulatory cytokines interleukin (IL)-10 and IL-27 ^2^. In humans, Tr1 cells have been identified in antigen-driven chronic inflammatory conditions, such as allogeneic hematopoietic stem cell transplantation (allo-HSCT)^3-5^, allergy (to bee venom, pollen, dust mites)^6-8^, autoimmunity (celiac disease^9^), infections (leishmania, malaria, influenza)^10-13^, as well as in cancer^14,15^. However, transcription factors (TFs) that control human Tr1 cell differentiation, phenotype and functions are largely unknown^2^. Tr1 cells do not express or depend on TF FOXP3, which determines the differentiation and function of thymic (t)Tregs^16^, and bestows the Treg-like phenotype and functions to FOXP3^+^ inducible (i)Tregs^17-19^. Tr1 lineage-determining TFs have been described in the murine, IL-27-inducible Tr1 differentiation model^20-24^, which has not yet been recapitulated in human cells^25^. Thus, the identification of TFs that govern human Tr1 cells requires the use of human Tr1 cell differentiation model. TFs EOMES^26^ or interferon regulatory factor 4 (IRF4) acting together with aryl hydrocarbon receptor (AHR)^27^ have been proposed as key regulators of Tr1 cells, but no consensus has been reached on the unifying TF signature that defines the Tr1 cell identity. In addition, EOMES and IRF4 and AHR were validated in polyclonal, not antigen-specific Tr1 cells, even though studies in murine disease models and human subjects suggest that Tr1 cell differentiation is antigen-driven^3,5-9,14,23,28^. Moreover, antigen-driven Tr1 cell differentiation can be recreated *in vitro* to generate clinically relevant Tr1 cell products, designed to induce antigen-specific tolerance^29,30^. Thus, revealing the key TFs of human antigen-specific Tr1 cells is critical to understand Tr1 biology, identify Tr1 cells *in vivo* and to provide a mechanistic basis for rational design of antigen-specific Tr1 cell therapies.

T-allo10^29,30^ is a human alloantigen-inducible Tr1 differentiation model designed to recreate the *in vivo* Tr1 induction observed after allo-HSCT, the condition where Tr1 cells were first identified^3^. During T-allo10 culture, allogeneic tolerogenic dendritic cells stimulate the alloreactive fraction of peripheral CD4^+^ Teff to clonally expand and differentiate into Tr1 cells^29^. The resulting Tr1 cells are oligoclonal and alloantigen-specific; namely, they selectively suppress alloreactive Teff, but not Teff activated by irrelevant antigens^29^. This alloantigen-specific suppressive activity makes T-allo10 Tr1 cells an attractive therapy to induce tolerance after allo-HSCT, as they could prevent graft-vs-host disease (GvHD) mediated by alloreactive Teff, without impairing Teff responses against tumor cells or pathogens^4,29,31^. HSCT donor-derived T-allo10 Tr1 products are already being tested in two phase I/Ib clinical trials (ClinicalTrials.gov ID NCT03198234 and NCT04640987), with promising early clinical results^32,33^. Using this clinically relevant Tr1 differentiation model, we recently compared the bulk transcriptome of human alloantigen-specific Tr1 cells to other CD4^+^ T cell subsets^29^. These data revealed that the TF IRF4, but not AHR or EOMES are significantly upregulated in human alloantigen-specific Tr1 cells^29^.

Inference of gene regulation from combined profiling of the epigenome and the transcriptome has been successful in identifying the transcriptional regulators of T cell differentiation and function^34^. Herein, we combined profiling of the epigenome and the transcriptome to identify the landscape of transcriptional regulators of human Tr1 cells. Single-cell transcriptomic, epigenomic, and functional genomic analyses of antigen-specific Tr1 cells induced *in vitro* revealed a signature of TFs unique to Tr1 cells, with IRF4, BATF (basic leucine zipper ATF-like transcription factor) and MAF (MAF bZIP transcription factor) acting as key transcriptional regulators of Tr1 cells. This Tr1 cell TF signature identified Tr1 cells *in vivo*, in both peripheral blood and tissue.

## Results

### *In vitro*-induced human antigen-specific Tr1 cells are distinct from FOXP3^+^ Tregs and other CD4^+^ T cell populations on a single-cell level

Previously, we used bulk RNA-sequencing (-seq) to reveal that *in vitro*-induced human alloantigen-specific Tr1 cells have a gene expression signature enriched for co-inhibitory genes, including LAG3, CTLA4, PDCD1 (encoding PD-1), HAVCR2 (encoding Tim-3), and ENTPD1 (encoding CD39), among others^29^. Here, we further define the Tr1 gene expression signature at the single cell level, and compare it to FOXP3^+^ Tregs and conventional CD4^+^ T cells present in the T-allo10 product. Towards this goal, we generated two T-allo10 cell products enriched with alloantigen-specific Tr1 cells, using a 10-day co-culture of total peripheral CD4^+^ T cells with allogeneic tolerogenic dendritic cells as described^29^ (**Fig.S1A**). An aliquot of T-allo10 cells at day 9 of T-allo10 culture was purified for live CD4^+^CD3^+^ T cells prior to single-cell capture and 5’ single-cell (sc) RNA- and TCR-seq (**Table S1**). The expected Tr1 cell phenotype and function was confirmed in the T-allo10 cell products at the end (day 10) of the T-allo10 culture, by measuring co-expression of surface proteins CD49b and LAG3 and anergy (**Fig.S1B-D**).

T-allo10 cells separated into 7 clusters (**Fig.1A**), which we annotated using data-driven and knowledge-based methods (**Fig.1B, Fig.S2A-C**). Except cluster C6, which was predominantly found in one donor, all other clusters were proportionally distributed (**Fig.S2D**). The proportion of Tr1 cells identified by sc-RNA-seq was similar to the proportion of Tr1 cells identified by flow cytometry (**Fig.S2D, Fig. S1C**). As expected from antigen-induced cells^29^, the Tr1 cluster contained the majority of clonally expanded cells (**Fig.1A**, **Fig.S2E**); the other cluster with highest number of clones was the small Th1 cluster C6 (**Fig.S2E**). Overall, in T-allo10 products, we detected clusters of FOXP3^+^ Tregs, naïve, resting, and polarized T cells that were distinguished from the Tr1 cell cluster (**Fig.1A**). Even using stringent filtering (FDR-adjusted *p*-value < 0.001 and absolute log2 fold-change ≥ 1), FOXP3^+^ Tregs and Tr1 cells had 49 differentially expressed genes (**Fig.1C, Table S2**). FOXP3^+^ Tregs expressed high level of FOXP3, IL2RA (encoding CD25), TIGIT, CD27, and TF BATF, while Tr1 cells expressed high levels of LAG3, interferon (IFN)-regulated IFIT gene family (IFITM1, IFITM2, IFITM3), FURIN (a protease which can activate latent transforming growth factor-β), genes conferring the cytotoxic phenotype (KLRB1, GZMA), and TF BHLHE40, which regulates IFN-γ and IL-10 production in Tr1 cells^35^ (**Fig.1C**).

**Figure 1.**
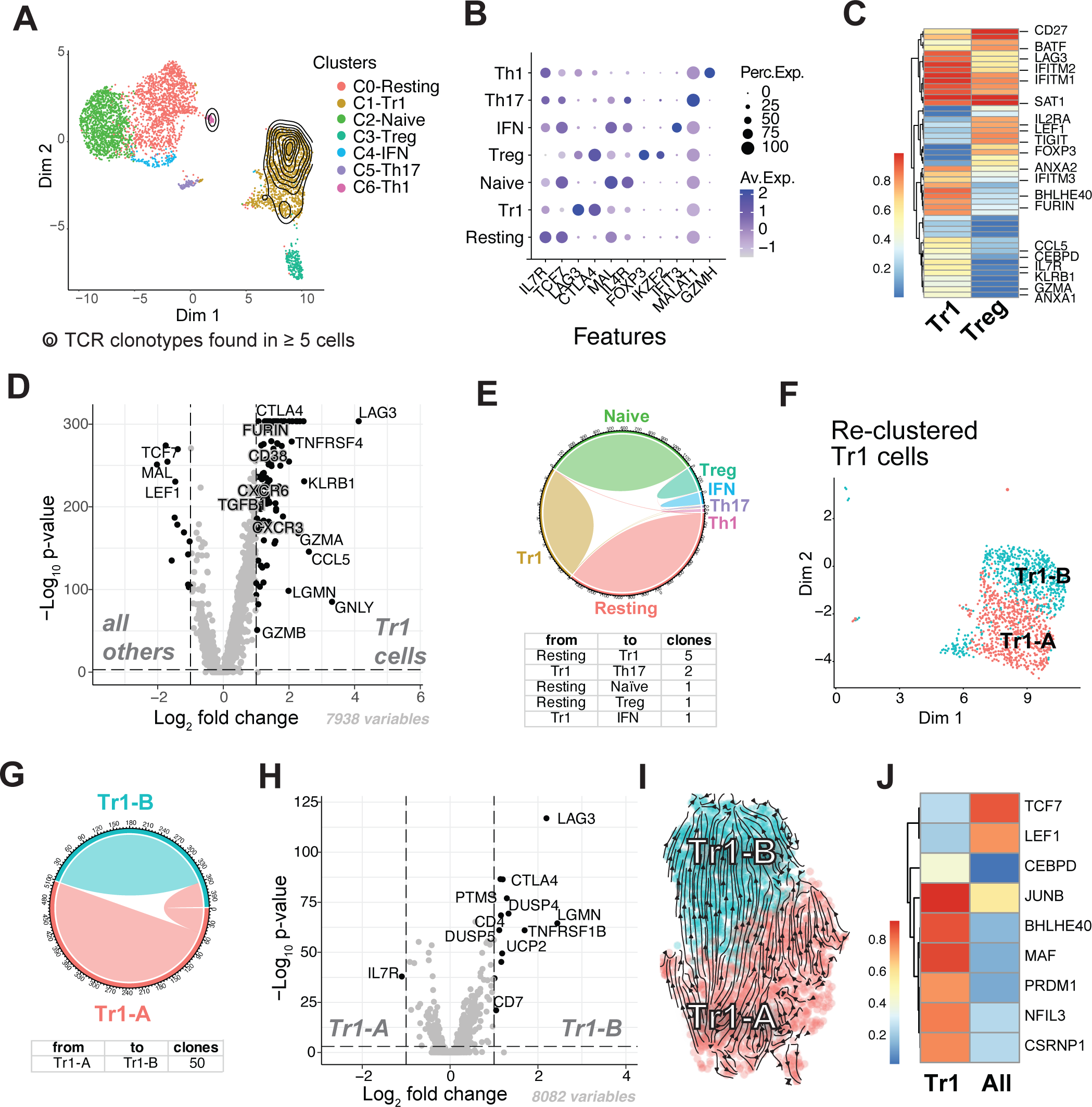
Single-cell RNA+TCR-seq analysis identifies antigen-specific Tr1 cells as a distinct population within T-allo10 cells. **A.** UMAP visualization of CD4^+^ T cells subpopulations within T-allo10 cells (n = 2). Overlaid contour plot shows clonally expanded cells. The lowest clonal frequency displayed is 5 cells. Dim = dimension. **B.** Dot plot illustrating the representative differentially expressed genes (DEG) per cluster. Dot color intensity corresponds to the average expression level per cluster. Dot size represents the percentage of cells expressing the gene within the cluster. **C.** Heatmap of average gene expression of 49 differentially expressed genes identified between Tr1 and FOXP3^+^ Treg cell clusters. Adjusted *p* (padj) value ≤ 0.001 and log2 fold change (log2FC) ≥ 1. **D.** Volcano plot depicting the results of the differential gene expression analysis between Tr1 and all other clusters. DEG that passed the filtering (*p*adj ≤ 0.001, log2FC ≥ 1) are in black; representative genes are labeled. **E.** Top: chord diagram showing clonotypes shared between clusters. Each arc represents a UMAP cluster from panel A, and ribbons connecting arcs represent shared clonotypes. The width of the ribbons corresponds to the number of shared clonotypes. Bottom: summary table provides the count of shared clonotypes between clusters. **F.** UMAP projection displaying the re-clustering of Tr1 cell cluster from A. **G.** Chord diagram showing the total number of unique and shared clones among Tr1 sub-clusters. **H.** Volcano plot illustrating the differential gene expression analysis among subclusters of Tr1 cells. DEG (*p*adj ≤ 0.001 and log2FC ≥ 1) are in black**. I.** Developmental trajectory of Tr1 cells predicted by RNA velocity. **J.** Heatmap representing the average gene expression of 9 differentially expressed transcription factors between Tr1 cells and the remaining clusters.

A comparison of Tr1 cell transcriptome to all other clusters revealed a distinct gene expression signature comprised from 90 significantly upregulated genes in Tr1 cells, including LAG3, CTLA4, FURIN, TGFB1, and cytotoxicity-associated genes KLRB1, GZMA, GZMB, GNLY and CCL5 (**Fig.1D, Table S3**). One quarter of these genes was also found in our published signature of alloantigen-specific Tr1 cells, derived from bulk RNA-seq analysis of Tr1 cells isolated from T-allo10 cell products based on co-expression of CD49b^+^ and LAG3^29^(**Fig.S3A**). We observed low expression of EOMES (**Fig.S3B**), which was previously described as a marker of polyclonal Tr1 cells and tissue Tr1 cells^14,26,36^, and a positive regulator of proteins associated with cytotoxic function (perforin, granzyme B, IFN-γ) in cytotoxic T cells^37^. Notably, despite the low expression of EOMES, Tr1 cells express granzyme B protein (encoded by GZMB) and are functionally cytotoxic (**Fig.S3C-D**).

The TCR repertoire of *in vitro*-induced antigen-specific Tr1 cells was oligoclonal and distinct from the repertoire of FOXP3^+^ Tregs and other CD4^+^ T cell populations in T-allo10 cell products. Tr1 cells shared only 8 clones with resting, Th17-like, and IFN-exhibiting (IFN) T cells (**Fig.1E**). Overall, clonal sharing between T cell populations in the T-allo10 cell product was minimal (**Fig.1E**), likely due to the antigen stimulation in Tr1-polarizing conditions and clonally diverse parental CD4^+^ T cells^29^. Notably, FOXP3^+^ Tregs did not share any clones with Tr1 cells (**Fig.1E**) nor did they clonally expand in response to alloantigen stimulus (**Fig.1A**). These data suggest that the frequency of alloreactive FOXP3^+^ Tregs in peripheral blood CD4^+^ T cells is low, or that the starting Treg population in culture was too small to detect rare alloreactive FOXP3^+^ Tregs^38^.

Next, we computationally selected Tr1 cells and re-clustered them, revealing two sub-clusters: Tr1-A and Tr1-B (**Fig.1F**). Tr1 sub-clusters shared 50 clones, indicating a common origin (**Fig.1G**). While there were only 14 DEG between the two clusters, Tr1 cells in cluster Tr1-B exhibited higher levels of activation-inducible genes such as LAG3, CTLA4, and dual-specificity phosphatases (DUSP4 and DUSP5), but lower IL7R (encoding the IL-7 receptor α-chain) (**Fig.1H, Table S4**). RNA velocity analysis^39^ suggested that the less activated Tr1 phenotype in cluster Tr1-A precedes more activated Tr1 cell phenotype in cluster Tr1-B (**Fig.1I**). Only one TF, JUNB, was differentially expressed between two clusters (2.24-fold higher in Tr1-B; adjusted *p*-value = 6.24 × 10^-46^). Since JUNB is downstream of T cell receptor (TCR) signaling, its expression in the Tr1-B cluster is in accordance with the more activated state of Tr1-B cells.

Finally, to identify the TFs that were significantly upregulated in Tr1 cells but not in other T cell populations, we examined the differentially expressed TFs between Tr1 cells and all other clusters. These analyses revealed 7 specifically upregulated TFs in Tr1 cells: CEBPD, JUNB, BHLHE40, MAF, PRDM1, NFIL3, and CSRNP1 (**Fig.1J**). Among these TFs, the role of CEBPD and CSRNP1 in T cells is poorly understood. We previously identified BHLHE40 as specific for *ex vivo* isolated Tr1 cells, showing it regulates IL-10 and IFN-γ^35^. NFIL3 was shown to repress FOXP3^40^ and upregulate IL-10 and HAVCR2 (encoding Tim-3)^41^. Similarly, MAF and PRDM1 are known regulators of IL-10^42,43^, and have also been implicated as regulators of murine Tr1 cell differentiation^23,43,44^. Altogether, our single-cell RNA- and TCR-seq analysis uncovered that *in vitro*-induced antigen-specific Tr1 cells have a distinct transcriptional signature and a TCR repertoire, and express a unique TF profile.

### Human antigen-specific Tr1 cells have a unique signature of accessible chromatin peaks

To further investigate the transcriptional regulation of Tr1 cell identity and obtain a comprehensive landscape of TF activity in antigen-specific Tr1 cells, we applied Assay for Transposase-Accessible Chromatin using sequencing (ATAC-seq^45^) and RNA-seq. First, we generated five T-allo10 products and control T-allo products (which contain CD4^+^ Teff cells) from the same parental CD4^+^ T cells (**Fig.2A**). We isolated CD49b^+^LAG3^+^ Tr1 cell and CD49b^-^LAG3^-^ non-Tr1 cell (DN, double-negative) populations from memory CD4^+^ T cells of T-allo10 products, and CD49b^-^LAG3^-^ Teff cells from control T-allo products by FACS, thawed cryopreserved parental CD4^+^ T cells, and analyzed these populations by ATAC- and RNA-seq. After quality control (**Fig.2B; Fig.S4A-B**), we compared the chromatin accessibility of Tr1 cells vs. all control populations combined, revealing 881 significantly accessible peaks in Tr1 (**Fig.2C**). Next, we annotated the peaks based on their nearest gene, and analyzed them using Gene Set Enrichment Analysis (GSEA)^46^, using Molecular Signature Database (MsigDB) gene set C7: Immune Signatures^47^. Data show that the genes that are significantly accessible in Tr1 cells were enriched in the gene signatures of regulatory T cells and memory T cells comprised in the MsigDB (**Fig.2D**). These signatures were also identified in Tr1 cells in our published, bulk RNA-seq dataset^29^, showing reproducibility across samples and platforms. Finally, we used Ingenuity Pathway Analysis (IPA)^48^ Upstream Regulator analysis to predict key transcriptional regulators of Tr1-specific accessible genes, identifying IRF4, MAF, and BHLHE40 TFs as top transcriptional regulators of Tr1 cells (**Fig.2E; Table S5**). These TFs were also significantly upregulated in Tr1 cells in either bulk^29^ or single-cell datasets or both (**Fig.S3A**). Chromatin peaks that were significantly accessible in Tr1 cells compared to control populations overlapped with known IRF4 and MAF promoter and enhancer regions (**Fig.2F**). In summary, these data show that the differentiation of antigen-specific human Tr1 cells leads to significant chromatin remodeling, with a distinct pattern of accessible chromatin peaks observed in Tr1 cells. Network enrichment analysis of Tr1-specific accessible peaks identified IRF4, MAF, and BHLHE40 TFs as top upstream regulators.

**Figure 2.**
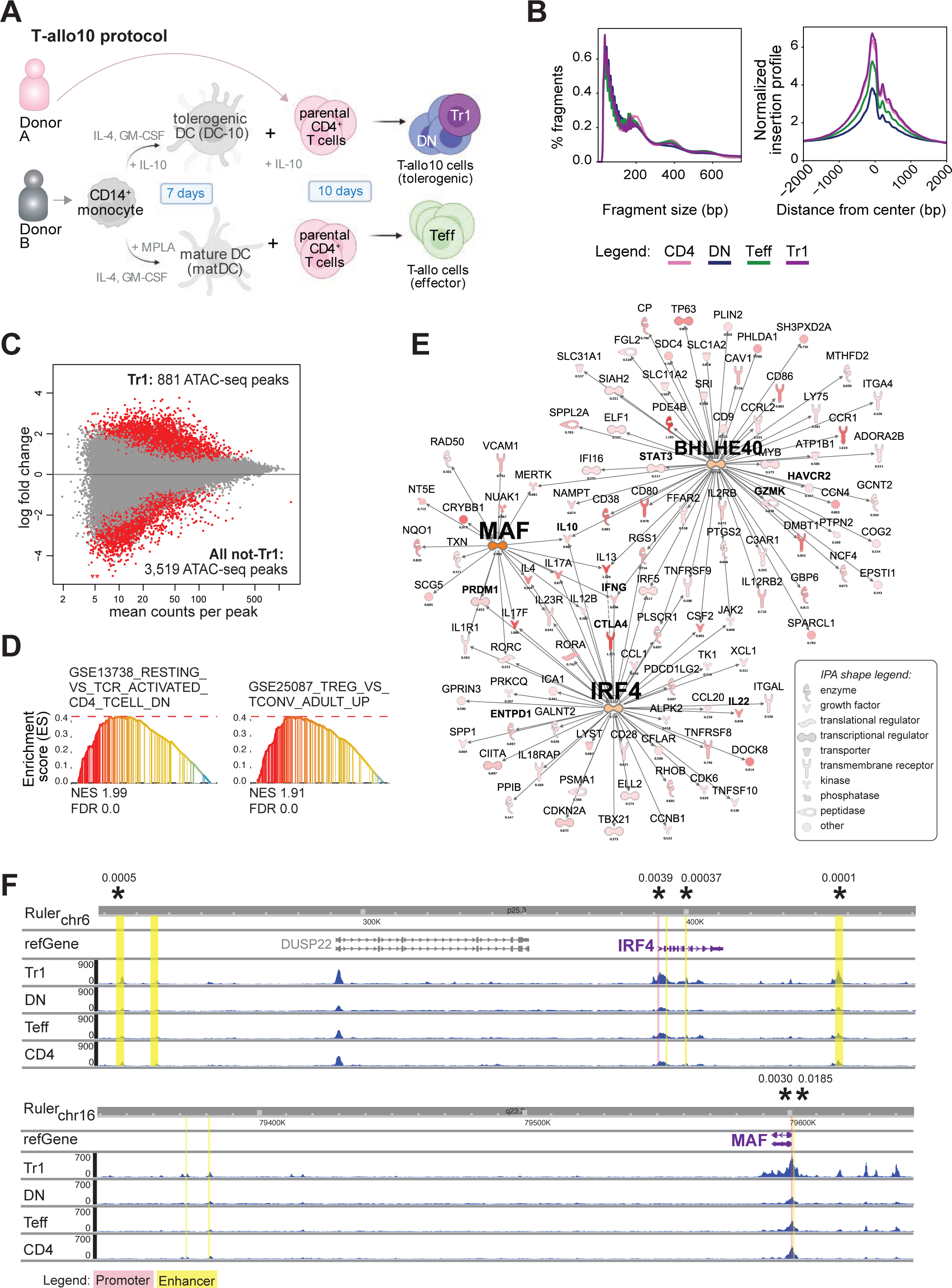
ATAC-seq reveals the chromatin accessibility landscape and transcriptional regulators of antigen-specific Tr1 cells. **A.** Experimental outline. Tr1 = CD49b^+^LAG3^+^ CD45RA^-^ CD4^+^ type 1 regulatory T cells from T-allo10 cell product; DN = double-negative (non-Tr1) CD49b^-^ LAG3^-^ CD45RA^-^CD4^+^ cells from T-allo10 product; Teff = CD49b^-^LAG3^-^ CD45RA^-^CD4^+^ effector T cells from control T-allo product; DC-10 = allogeneic tolerogenic dendritic cells; matDC = allogeneic mature dendritic cells; MPLA = monophosphoryl lipid A, a TLR4 agonist given at day 5 of matDC induction. **B.** Quality control analysis (QC) of bulk ATAC libraries showing nucleosome periodicity (left) and transcription start site enrichment (right). Data are averaged from n = 5 per condition. **C.** Changes in chromatin accessibility in Tr1 cells (n = 5) relative to all other non-Tr1 cells (CD4, Teff, DN, total n = 15). Each dot represents one ATAC-seq peak. **D.** Gene Set Enrichment Analysis (GSEA) of ATAC-seq peaks of Tr1 cells versus all non-Tr1 cells within MsigDB C7: Immune Signatures database; representative enrichment plots are shown. NES = normalized enrichment score; FDR = FDR *q* value. **E.** Upstream regulator Ingenuity Pathway (IPA) analysis of the top 2,336 accessible peaks in Tr1 cells, ranked by the number of times found differentially accessible between Tr1 cells and all other T cell subsets (≥ 0.7) and filtered on transcriptional regulators that were present in the dataset, showing the predicted top three most significant upstream regulators (hub centers) in a network with downstream genes. Color = log2FC. **F**. Gene accessibility track visualization of Tr1-specific transcription factors IRF4 and MAF. Promoter regions are highlighted in red, while enhancer regions are highlighted in yellow. Significantly accessible regions in Tr1 cells compared to the non-Tr1 cells are marked with asterisks (*), with corresponding *p*-value indicated numerically.

### Human antigen-specific Tr1 cells have a distinct TF usage profile

In addition to revealing global chromatin accessibility, ATAC-seq can also identify TFs that are actively used by the cell even if their gene expression is low. TF usage can be identified by i) mapping the known TF binding sites (“motifs”) to the areas of accessible chromatin, ii) mapping the TF footprints to identify TFs bound to the chromatin at the time of sampling, iii) uncovering the TFs that colocalize with predicted enhancer regions, and iv) integrating ATAC- and RNA-seq data to identify the TFs with high peak accessibility-gene expression associations. Active TFs in Tr1 cells that are identified by both peak accessibility and TF usage analyses are likely to be biologically relevant. We analyzed TF motif accessibility in Tr1 cells and control cell populations using ChromVar^49^, and showed that Tr1 cells cluster apart from other cell subsets (**Fig.3A**). The TF motifs that were significantly more accessible in Tr1 cells compared to control populations contained AP-1 (Fos, Jun and others), MAF, IRF (Interferon Regulatory Factor), and T-box TF family members, among others (**Fig.3B**). We also performed footprinting analysis using TOBIAS^50^, which was concordant with the motif analysis, revealing many of the same TF family members as significantly more active in Tr1 cells (**Fig.3C**). TFs with prominent footprints included IRF4, which we previously identified as upregulated in Tr1 cells^29^ and which others have found to be important in polyclonal murine^24^ and human^27^ Tr1 cell differentiation; the BATF:JUN heterodimer, which may facilitate IRF4 expression^51^; RUNX2, which was found in cytotoxic CD8^+^ T cells^52^ and NK cells^53^ but not previously described in Tr1 cells; and BHLHE40, which regulates IL-10 and IFN-γ production in *ex vivo* human Tr1 cells from peripheral blood^35^ (**Fig.3D**). TFs IRF4, MAF, PRDM1, TBX21 and RUNX2, for which the antibodies for flow cytometry were available, were also upregulated in Tr1 cells at the protein level (**Fig.3E**).

**Figure 3.**
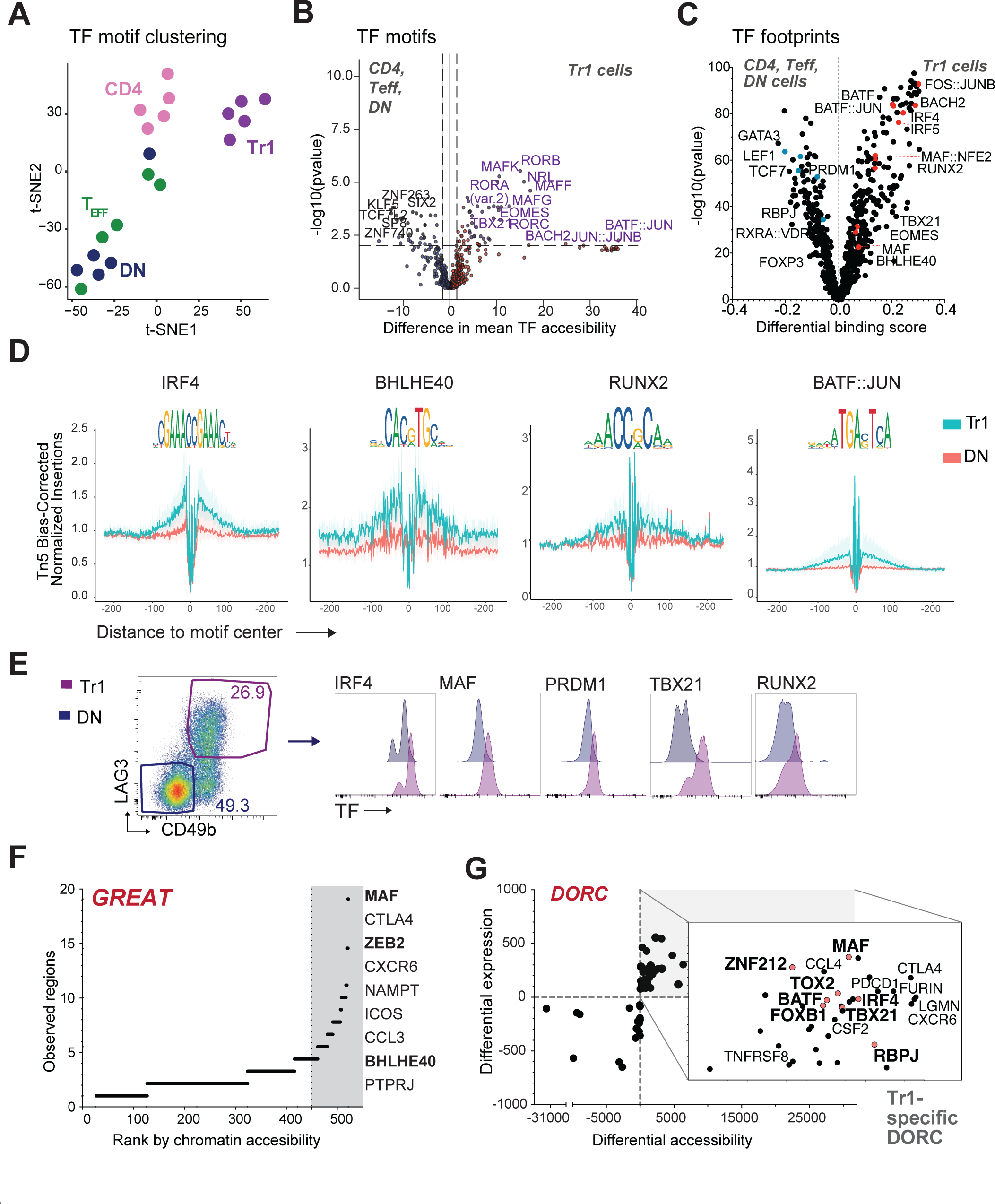
Antigen-specific Tr1 cells have a distinct signature of active transcription factors. **A.** tSNE visualization of ATAC-seq profiles using 452 transcription factor (TF) motif deviations identified by chromVAR package. Samples are colored by cell type. **B.** Volcano plot showing transcription factor enrichment (chromVAR) between Tr1 and all non-Tr1 cells. *P*-values were calculated using a two-tailed t-test. **C.** Volcano plot showing the differential binding activity, calculated by TOBIAS package, between Tr1 and all non-Tr1 cells; each dot represents one motif. Representative Tr1-specific TFs are highlighted in red, and non-Tr1 TFs in blue. **D.** Bias corrected ATAC-seq footprints between Tr1 and DN (non-Tr1) cells from T-allo10 products. Selected motifs that have differential binding activity between Tr1 and all non-Tr1 cells (panel C) are shown. **E.** Representative histograms of TF expression within Tr1 and non-Tr1 (DN) populations of T-allo10 cells. n = 2 - 6 per TF. **F.** Enhancers scores predicted by GREAT, ranked by chromatin accessibility. Representative ranked-enhancers among the top 50 are shown on the right. **G.** Paired correlation between differential expression and differential accessibility, computed using the Domains of Regulatory Chromatin (DORC) calling method from the FigR package. Highlighted top right quadrant shows the Tr1-specific DORC and top associated genes (normal font, black dots) and TFs (bold font, red dots).

Next, we asked which TFs co-localize with regions of the Tr1 genome exhibiting super-enhancer-like features. Super-enhancers are large clusters of enhancers with high TF binding activity, associated with expression of TFs and genes crucial to cell identity^54,55^. GREAT algorithm extrapolates biological functions of non-coding genomic regions by querying the annotations of nearby genes, and can infer enhancers from ATAC-seq data^56^. Highest ranked enhancers, which might represent super-enhancers, were found nearby TFs MAF, BHLHE40, and ZEB2, and archetypal Tr1 genes^29^ such as CTLA4, NAMPT and CCL3 (**Fig.3F, Table S6**). Interestingly, ZEB2 was recently shown to induce GZMA (encoding granzyme A)^57^, which is highly expressed in *in vitro*-induced antigen-specific Tr1 cells (^29^, **Fig.1E, Fig.S2C, Table S3**).

As a final step, we integrated ATAC-seq data with matched RNA-seq data (**Fig.2A**) using FigR^58^, which identified domains of regulatory chromatin (DORC). DORC are chromatin regions with high-density associations between cis-regulatory elements and genes, which are enriched for lineage-determining genes and overlap with known super-enhancers^58,59^. The Tr1 cell-specific DORC (**Table S7**) contained archetypal Tr1 genes such as CTLA4, PDCD1, and FURIN (**Fig.3G, Fig.S4C, Table S3**), and the following TFs: MAF, IRF4, BATF, TBX21 (encoding T-bet), TOX2, RBPJ, FOXB1 and ZNF212 (**Fig.3G**, **Fig.S4C**). Altogether, our comprehensive bioinformatic analysis of TF usage show that antigen-specific human Tr1 cells have a unique profile of active TFs, and identified MAF and IRF4 as one of the top candidate TFs that regulate the key biological features of Tr1 cells.

### IRF4, BATF and MAF regulate human antigen-specific Tr1 cells

Next, we investigated the biological relevance of top candidate TFs using functional genomics. We started with IRF4 and MAF, which were identified as top upstream regulators of Tr1 cells in network enrichment analysis (**Fig.2F**), in TF usage analysis (**Fig.3B-C**), in Tr1 DORC (**Fig. 3G**), and they were highly expressed in Tr1 cells (^29^, **Fig.1J).** BATF expression was higher in FOXP3^+^ Tregs than in Tr1 cells (**Fig.1C**), but we also validated its role because BATF is often co-regulated with IRF4^51,60,61^, and was identified in Tr1 cells in the TF usage and DORC analyses (**Fig.3B-C** and **Fig.3G**, respectively). Our first approach was to knock-out (KO) the target TFs in parental CD4^+^ T cells using CRISPR/Cas9 before Tr1 cell differentiation in T-allo10 culture (**Fig.4A**). The efficiency of CRISPR-based gene editing efficiency in primary T cells is usually increased by TCR pre-stimulation, but this strategy prevented antigen-inducible Tr1 cell differentiation in T-allo10 culture (our unpublished data). Instead, we knock-out the TFs from unstimulated, *ex vivo* isolated CD4^+^ T cells with multiplex single guide (sg) RNAs, designed to disrupt the universal second exon of IRF4 and first exons of BATF and MAF (**Fig.S5A**). Multiplex sgRNA strategy achieved good KO efficiencies in unstimulated CD4^+^ T cells (> 75% median; **Fig.4B**).

**Figure 4.**
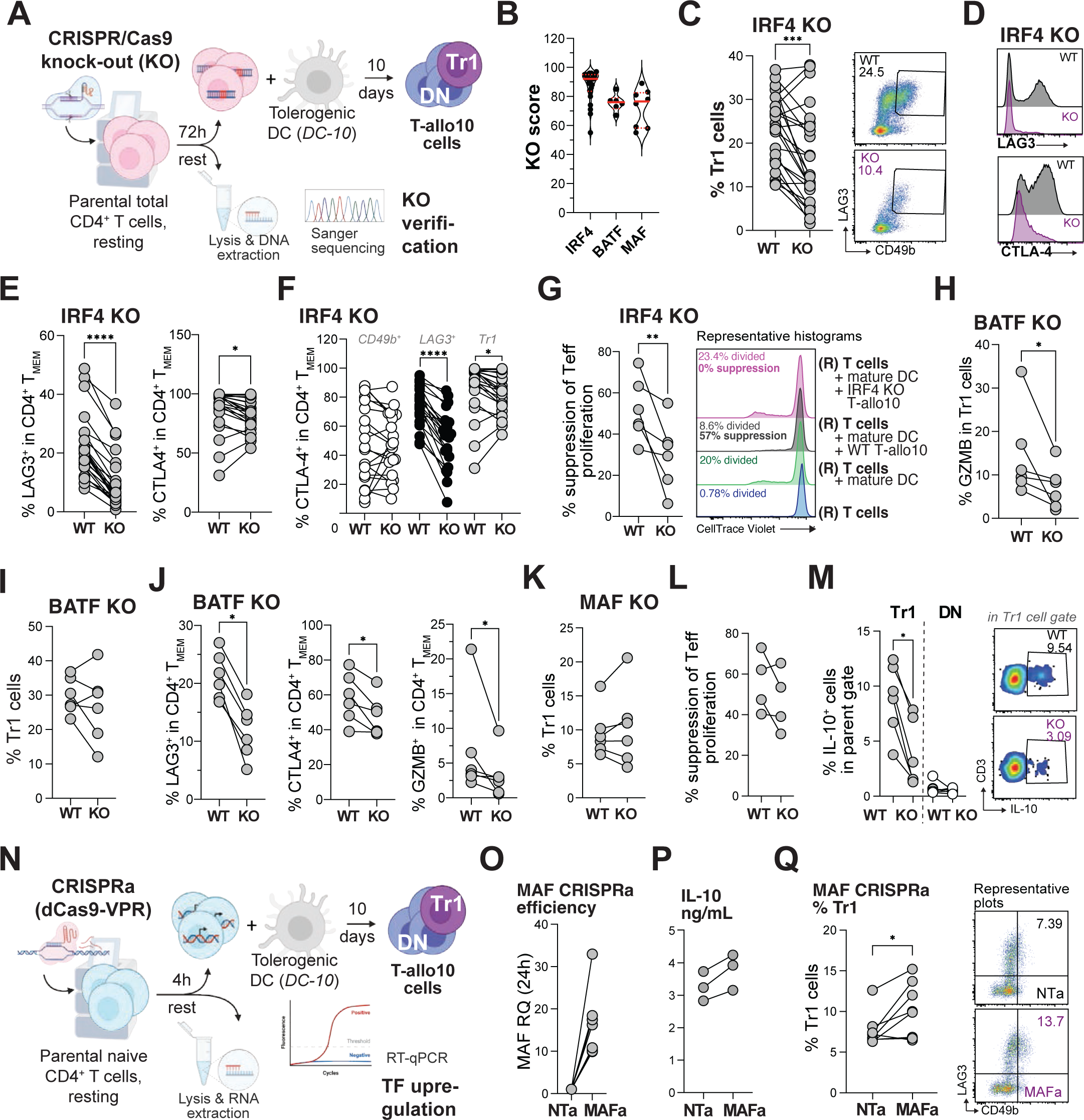
Transcription factors IRF4, BATF and MAF play non-redundant roles in Tr1 cell differentiation, phenotype, and function. **A.** Strategy for the transcription factor knock-out (KO) in parental CD4^+^ T cells, combined with their differentiation into antigen-specific Tr1 cells. DC = dendritic cells. **B.** Knock-out score (percent) in CD4^+^ T cells 72h post-editing, as calculated by ICE (Inference of CRISPR Edits, Synthego) using Sanger sequencing data. Each dot represents a healthy donor sample, red full lines the median, and red dotted lines interquartile range. IRF4 KO, n = 25; BATF KO, n = 6; MAF KO, n = 8. **C.** Left: cumulative flow cytometry data of Tr1 cell frequency at the end of 10-day T-allo10 culture, started from non-targeted (wild-type, WT) or IRF4 KO parental CD4^+^ T cells that were co-cultured in 10:1 ratio with allogeneic DC-10; n = 25. Right: representative flow cytometry plots showing Tr1 cell staining in WT and IRF4 KO memory CD4^+^ T cells within T-allo10 products. Numbers on plot indicate frequency of gated cells. **D.** Representative flow cytometry histograms depicting the expression of LAG3 and CTLA-4 in memory CD4^+^ T cells of WT and IRF4 KO T-alo10 products. **E.** Frequency of LAG3^+^, CD49b^-^ cells (left), and CTLA-4^+^ cells (right) in CD4^+^ memory T cells (Tmem) in WT or IRF4 KO T-allo10 cells; n = 25, flow cytometry **F.** Frequency of CTLA-4^+^ cells in CD49^+^LAG3^-^ (white circles), CD49b^-^ LAG3^+^ (black circles), and CD49b^+^LAG3^+^ Tr1 cells (grey circles), within Tmem of T-allo10 cells; n = 25, flow cytometry. **G.** Left: Suppression of responder (“R”) T cell proliferation by WT or IRF4 KO T-allo10 cells. R = *ex vivo* isolated CD4+ T cells autologous to T-allo10 cell donors. Suppression assay was performed as described^29^. Right: representative histograms of dividing responder (“R”) CD4+ T cells labeled with CellTrace Violet (CTV) proliferation dye, cultured for 5 days alone (blue), or stimulated with allogeneic mature dendritic cells (DC) ± WT or KO T-allo10 cells. Flow cytometry, n = 6. **H.** Frequency of granzyme B (GZMB)^+^ cells in Tr1 cells from WT or BATF KO T-allo10 cells; n = 6, flow cytometry. **I.** Frequency of Tr1 cells in WT or BATF KO T-allo10 cells (as in C); n = 6, flow cytometry. **J.** Frequency of LAG3^+^, CD49b^-^ cells (left), CTLA-4^+^ cells (middle), and granzyme B (GZMB)^+^ cells in CD4^+^ memory T cells (Tmem) from WT or BATF KO T-allo10 cells; n = 6, flow cytometry. **K.** Frequency of Tr1 cells in WT or MAF KO T-allo10 cells (as in C); n = 6, flow cytometry. **L.** Suppression of responder (“R”) T cell proliferation by WT or MAF KO T-allo10 cells (as in G); n = 4, flow cytometry. **M.** Frequency of IL-10^+^ CD49b^+^LAG3^+^ Tr1 cells or non-Tr1 (CD49b^-^LAG3^-^ DN or double-negative) cells in Tmem of WT or MAF KO T-allo10 products. Left: cumulative data, n = 5; right: representative dot plots, flow cytometry. **N.** Strategy for the transcription factor (TF) overexpression in parental naive CD4+ T cells using CRISPRa (CRISPR-activation), combined with their differentiation into Tr1 cells. dCas9-VPR = “dead”, nuclease-inert Cas9 fused to transcriptional activation domains VP64, p65 and Rta; RT-qPCR = real-time quantitative PCR. **O.** CRISPRa efficiency, measured by upregulation of MAF gene 24h after CRISPRa (MAFa) relative to non-targeted controls (NT) using RT-qPCR; n = 8. **P.** Frequency of Tr1 cells after MAF CRISPRa in comparison to NT in Tmem cells of T-allo10 products. Left: cumulative data; right: representative dot plots. * *p* < 0.05, *** *p* < 0.001, **** *p* < 0.0001; Wilcoxon test.

After 10 days of *in vitro* differentiation, T-allo10 cell products generated from KO and mock-treated wild-type (WT) parental CD4^+^ T cells were compared for Tr1 cell frequency, phenotype, and function. Compared to controls, IRF4 KO significantly decreased the yield of Tr1 cells, identified by the co-expression of CD49b and LAG3, within the T-allo10 cell products (**Fig.4C**). After IRF4 KO, the expression of LAG3 and CTLA-4 was also significantly decreased in memory CD4^+^ T cells (**Fig.4D-E**), while CTLA-4 expression was significantly decreased in LAG3-single-positive memory CD4^+^ T cells and Tr1 cells (**Fig.4F**). IRF4 KO did not affect the expression of granzyme B (**Fig.S5B**). Interestingly, the remaining WT CD4^+^ T cells outgrew the IRF4-KO CD4^+^ T cells in the Tr1 compartment, but not in non-Tr1 memory or naïve CD4^+^ T cells in IRF4-KO T-allo10 products, (**Fig.S5C**), suggesting that IRF4 provides a competitive growth advantage to *in vitro* differentiating Tr1 cells. Finally, we observed a significant impairment of the suppressive function of IRF4 KO T-allo10 cells (**Fig.4G**), indicating that IRF4 regulates suppressive capacity of Tr1 cells.

On the other hand, BATF KO significantly impaired the expression of granzyme B in Tr1 cells (**Fig.4H)**, but it did not affect Tr1 cell differentiation (**Fig.4I**) and CTLA4 expression in Tr1 cells (**Fig.4I Fig.S5D**). Interestingly, BATF KO impaired the ability of memory CD4^+^ cells to express LAG3, CTLA-4 and granzyme B (**Fig.4I and Fig.S5D**). We also tested the effect of MAF KO on Tr1 cell differentiation, phenotype, and function. MAF KO did not modulate Tr1 cell differentiation (**Fig.4K**), the expression of LAG3 or CTLA-4 (**Fig.S5E**), or suppressive function of T-allo10 cells (**Fig.4L**). However, in accordance with its described role in regulating IL-10, MAF KO significantly impaired the production of IL-10 in Tr1 cells (**Fig.4M**).

Because IL-10 drives the differentiation of Tr1 cells, we asked whether MAF overexpression (OE) can increase Tr1 cell yields *in vitro*. We designed sgRNA guides to the MAF promoter region and upregulated MAF in naive CD4^+^ T cells using CRISPR-activation (CRISPRa), which induces transient (3-5 days) OE of the target gene^62,63^ (**Fig.4N**). We achieved strong upregulation of MAF 24h after CRISPRa (**Fig.4O**). MAF OE induced a trend towards an increase in soluble IL-10 at the end of 10-day T-allo10 culture compared to non-targeted controls (**Fig.4P**) and significantly increased Tr1 cell yield at the end of the T-allo10 culture (**Fig.4Q**).

In summary, we show that IRF4, BATF, and MAF, identified by the bioinformatic analyses as top transcriptional regulators of *in vitro*-induced antigen-specific Tr1 cells, play key roles in regulating various aspects of Tr1 cell biology: their differentiation, phenotype, and functions.

### Tr1-specific TF signature is detected *in vivo* in peripheral blood CD4^+^ T cells

Next, we sought to understand if the Tr1 TF signature, which we identified in *in vitro* induced antigen-specific Tr1 cells, can be used to detect Tr1 cells *in vivo.* To this goal we applied CIBERSORTx, a digital cytometry method that estimates abundances of target cell types in a heterogenous bulk population using high-throughput data^64^. We analyzed the Tr1 cell frequencies by CIBERSORTx in ATAC-seq data of bulk CD4^+^ T cells isolated from peripheral blood of three patients with hematological malignancies, at three time points post-HSCT and treatment with donor-derived T-allo10 cells (ClinicalTrials.gov ID: NCT03198234;^29^; **Fig.S6A**). TF motif signatures of healthy donor CD4^+^ T cells, which contain significantly fewer CD49^+^LAG3^+^ Tr1 cells than T-allo10 recipients^29^ (**Fig.S6B-C**), were included to test the sensitivity of the method. We used the interactive CIBERSORTx user interface^64^ and previously identified TF motif signatures of purified parental CD4^+^ T cells and Tr1 and non-Tr1 cells from the T-allo10 cell products (**Fig.3A-B**) to construct a CIBERSORTx signature matrix (**Table S8**). Next, we investigated the relative accessibility of TF motifs, identified by the CIBERSORT signature matrix as specific for Tr1 cells, within the bulk CD4^+^ T cells. Data show that all but one patient sample clustered apart from healthy donor samples (**Fig.5A**). Based on this TF matrix and TF motif accessibility values, CIBERSORTx determined the Tr1 cell frequencies in bulk CD4^+^ T cell samples which we validated by comparing them to Tr1 cell frequencies obtained by flow cytometry. These data revealed that the frequencies of Tr1 cells in the three patients at all time points were significantly higher than the frequencies observed in the healthy donors (**Fig.S6-C**), and that the frequency of Tr1 cells identified by digital cytometry significantly correlated with the frequency of Tr1 cells identified by flow cytometry (**Fig.5B**). These data confirm our previous TCR clonal tracking data that Tr1 cells persist long-term *in vivo* after adoptive transfer^29^, and demonstrate that the Tr1 TF signature identified in *in-vitro* induced antigen-specific Tr1 cells can be leveraged to detect Tr1 cells *in vivo*.

**Figure 5.**
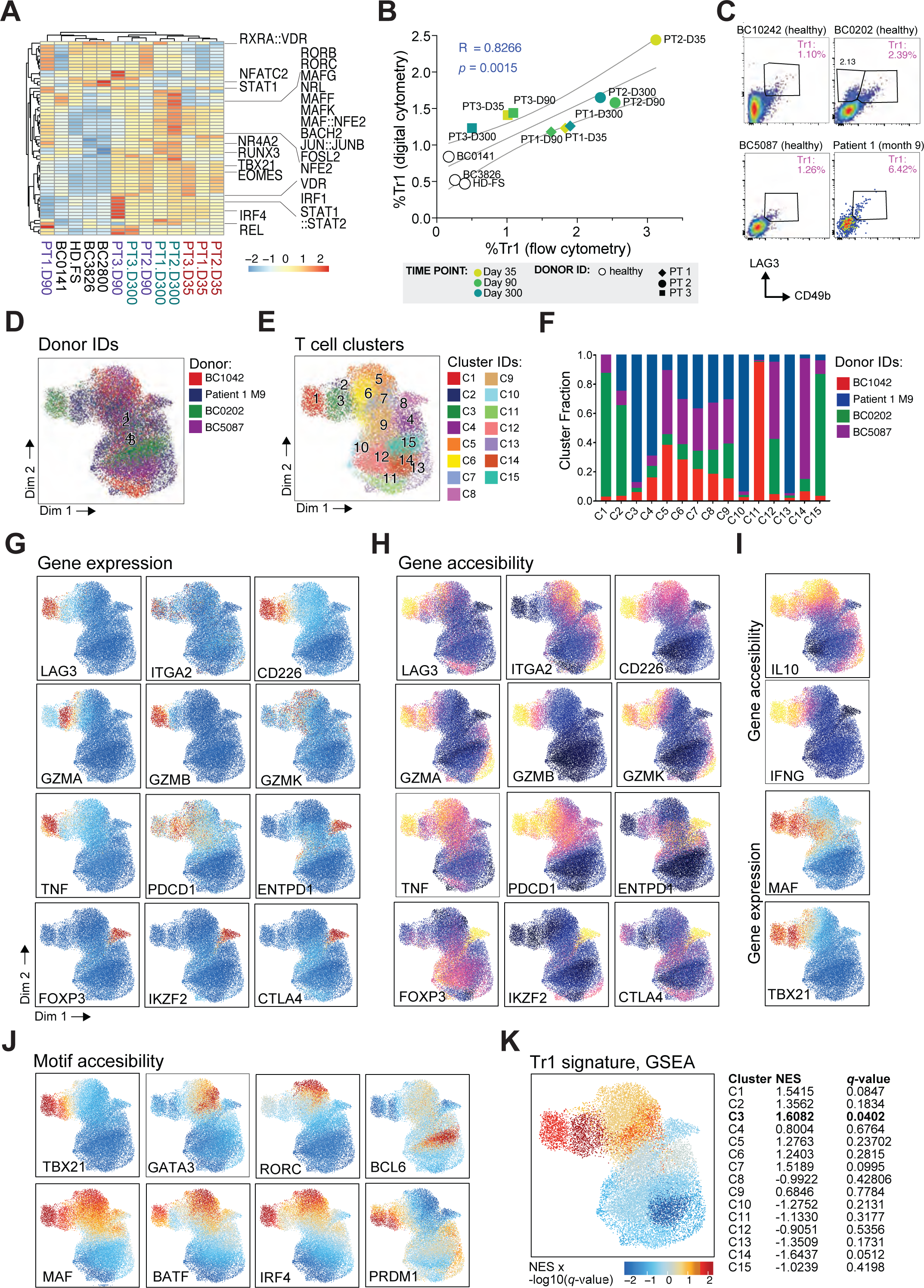
Identification of peripheral blood Tr1 cells *in vivo* using Tr1-specific transcription factor signature. **A.** Heatmap of transcription factor (TF) motifs identified as specific to T-allo10-derived Tr1 cells by CIBERSORTx algorithm, as expressed in total CD4^+^ T cells isolated from peripheral blood mononuclear cells (PBMC). HD and BC = healthy donors; PT = patients 1-3; D35, 90 and 300 indicate days post-allo-HSCT and T-allo10 therapy. **B.** Frequency of Tr1 cells estimated by digital cytometry (CIBERSORTx; y-axis) and measured by flow cytometry (x-axis) in total CD4^+^ T cells; Spearman rank correlation. Line indicates linear regression trendline. **C.** Frequency of Tr1 cells in total CD4^+^ T cells of healthy donors (n = 3) and a patient at month 9 post-allo-HSCT and T-allo10 infusion, which were subsequently analyzed by sc-multiome (single-cell ATAC+RNA-seq, 10X Genomics). **D.** Donor IDs after integration of samples with Harmony; ArchR package. **E.** sc-multiome clusters of CD4^+^ T cell samples from panel C; ArchR package. **F.** Distribution of donors across clusters. **G**. Gene expression of selected genes characteristic for Tr1 cells, cytotoxic T cells, and FOXP3^+^ Tregs. **H.** Gene accessibility of genes depicted in panel G. **I.** From top to bottom: gene accessibility of cytokines IL10 and IFNG and motif accessibility of TFs MAF and TBX21 that induce IL10 and IFNG, respectively. **J.** Motif accessibility of TFs that act as key regulators of T-helper cell subsets. **K.** Gene Set Enrichment Analysis (GSEA) of the accessible TF signature specific to *in vitro*-induced antigen-specific Tr1 cells in CD4^+^ T cells of healthy donors and a patient analyzed by sc-multiome. NES = normalized enrichment score; *q*-value = FDR-corrected *p*-value. Color scale = *q*-value-adjusted NES.

To confirm that the TF motif signature of Tr1 cells can also be used to identify Tr1 cells on a single cell level, we analyzed the purified peripheral blood CD4^+^ T cells from three healthy donors and one patient at 9 months post-T-allo10 infusion and allo-HSCT with sc-multiome (sc-ATAC + RNA-seq; 10X Genomics). All donors had detectable CD49b^+^LAG3^+^ Tr1 cells by flow cytometry (**Fig.5C**), with higher frequency of Tr1 cells in healthy donor BC0202 and patient 1 at month 9 (M9) than the other two healthy donors (**Fig.5C**). Samples were integrated, then clustered based on accessibility data (**Fig.5D**) because nuclear RNA was sparse in *ex vivo* isolated, unstimulated T cells (**Fig.S6D**). Clustering revealed 15 CD4^+^ T cell clusters (**Fig.5E, Table S9**), some of which were evenly distributed across donors, while others predominantly contained cells from one donor (**Fig.5F**). Clusters C1 and C3 were identified as putative Tr1 clusters based on the high gene expression of LAG3, ITGA2 (encoding CD49b) and CD226 (**Fig.5G**), which were previously shown to be specific to peripheral blood Tr1 cells^65^. These two clusters predominantly originated from healthy donor BC0202 (C1, **Fig.5F**) and the patient M9 (C3, **Fig.5F**). Clusters C1 and C3 expressed granzyme genes, but GZMB (encoding granzyme B) was higher in cluster C1, GZMA (encoding granzyme A) was higher in cluster C3, while GZMK was present in both (**Fig.5G**). Because GZMA expression is higher than GZMB in Tr1 cells (**Fig.1E**), we compared the gene expression between clusters C1 and C3. This comparison revealed that cluster C1 has significantly higher expression of GZMB, granulysin (GNLY), TNF (encoding TNF-α), LTA (lymphotoxin-α), several IFN-inducible genes from the IFIT and 2’-5’-oligoadenylate synthase (OAS) families, and TRBV28 gene (the variable region 28 of the TCR β-chain), among others (**Fig.S6E, Fig.5G**). On the other hand, cluster C3 contained more PDCD1 (PD-1)-expressing cells **(Fig.5G**), and higher level of archetypal Tr1 cell genes^29^ ENTPD1 (CD39), IRF4, and DUSP4 (**Fig.S6D, Fig.5G**). Thus, cluster C3 contains Tr1 cells, while cluster C1 likely contains a combination of Tr1 and Th1 or T_EMRA_^66^ (T effector memory cells re-expressing CD45RA) cells, given that ITGA2 (CD49b) is expressed by some cells in that cluster (**Fig.5G**). Interestingly, CTLA4 expression in Tr1 cell cluster C3 was low (**Fig.5G**). This may be due to sparsity of nuclear RNA data or a less activated state of these Tr1 cells *in vivo*. Indeed, we observed low CTLA4 expression in CD49b^+^LAG3^+^ Tr1 cells isolated from healthy donor peripheral blood by both bulk RNA-seq^35^ and on protein level, albeit still significantly higher than in non-Tr1 memory CD4^+^ T cells (**Fig.S6F**). As in T-allo10 products (**Fig.1**), FOXP3^+^ Treg cluster, C8, was clearly identified by high expression of FOXP3, Helios (encoded by IKZF2), and CTLA4 (**Fig.5G**).

Next, we analyzed accessibility of the same genes as in **Fig.5G**. Accessibility of LAG3 and CD226 or granzymes was not exclusive to Tr1 cell cluster C3 (**Fig.5H**). In contrast to its RNA expression pattern, ITGA2 gene was inaccessible (**Fig.5H**), (**Fig.5G**), suggesting a difference in regulation between chromatin accessibility and nuclear RNA transcription. The accessibility of CTLA4 in Tr1 cluster C3 was lower, but PDCD1 accessibility was higher than in FOXP3^+^ Tregs (**Fig.5H**). In addition, cluster C3 had accessible IL10 and IFNG genes, as well as their regulators MAF and TBX21, respectively (**Fig.5I**). Furthermore, we analyzed TF motif accessibility in this dataset, showing that Tr1 cell cluster C3 had accessible motifs of TBX21, MAF, BATF, IRF4 and PRDM1, among others (**Fig.5J**). None of these TF motifs were exclusively accessible in Tr1 cell clusters (**Fig.5J**). Notably however, Tr1 clusters did not have accessible GATA3, RORC, and BCL6 motifs, which are archetypal for Th2, Th17, and Tfh cells, respectively (**Fig.5J**).

To confirm our annotation of cluster C3 as the putative Tr1 cell cluster, we first derived a common signature of actively used TFs in *in vitro*-induced antigen-specific Tr1 cells (**Methods, Table S10**). Next, we used this Tr1-specific TF signature and GSEA to determine its enrichment in the sc-multiome dataset. Indeed, C3 was the only cluster that was significantly enriched for the Tr1-specific TF signature (**Fig.5K, Table S11**). This TF signature was not enriched in independent datasets of B cells and monocytes, confirming its Tr1 cell specificity (**Fig.S7A-B**). Altogether, we demonstrate that we can identify Tr1 cells in the peripheral blood of patients and healthy donors using a Tr1-specific TF signature.

### Tr1-specific TF signature identifies Tr1 cells in solid tumors

Next, we investigated if the Tr1-specific TF motif signature, which was identified in *in vitro*-induced antigen-specific Tr1 cells, can be broadly applied to identify Tr1 cells in tissues. To this aim, we re-analyzed published sc-ATAC-seq dataset from peripheral blood, tumor tissue and adiacent normal tissue of 8 patients with clear-cell renal cell carcinoma samples (ccRCC), 4 patients in stage 1A and 4 patients in stage 1 B^67^. We focused on CD4^+^ T cells while filtering out CD8^+^ T cells, B cells, NK cells, and myeloid cells (**Fig.6A; Methods**). By integrating the peripheral blood and tissue samples of 8 ccRCC patients (**Fig.6B)** we identified 12 CD4^+^ T cell clusters (**Fig.6C; Table S12**). Most clusters were found in all patient samples (**Fig.6D**) and proportionally distributed across patients with stage 1A or 1B ccRCC (**Fig.6E**). However, cells predominantly clustered together according to the tissue type (**Fig.6F-G**). GSEA analysis identified cluster C9, composed from naïve CD4^+^ T cells from peripheral blood, as significantly depleted from the Tr1 TF signature (**Fig.6H, Table S13**). Cluster C6, composed of tumor CD4^+^ T cells, was significantly enriched for Tr1 TF signature, with normalized enrichment score (NES) = 2.1371 (**Fig.6H, Table S13**). Accordingly, Tr1 cluster C6 had highly accessible motifs for TBX21, MAF, BATF and IRF4, but closed motifs characteristic for FOXP3^+^ Tregs, Th2, Th17 cells, and Tfh cells (FOXP3, GATA3, RORC, and BCL6, respectively) (**Fig.6I**).

**Figure 6.**
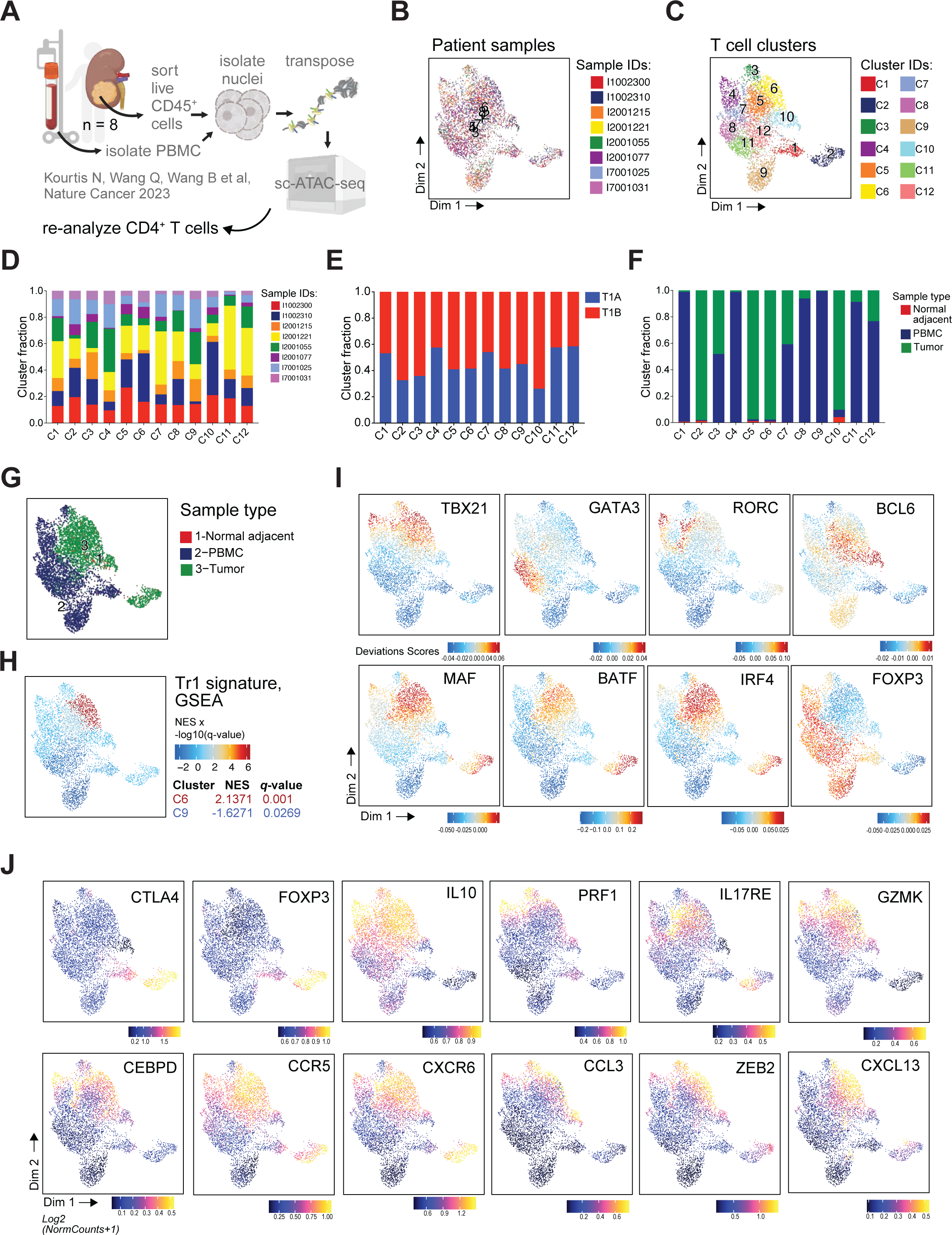
Identification of solid tumor-resident Tr1 cells *in vivo* using Tr1-specific transcription factor signature. **A.** Computational analysis workflow applied to publicly available single-cell ATAC-seq data from clear-cell renal cell carcinoma (ccRCC, n = 8)^67^. The 10x Genomics output was processed using ArchR to analyze CD4^+^ T cells after filtering out myeloid cells, NK cells, and B cells from total CD45^+^ cells from ccRCC samples. **B.** UMAP representation of 8 ccRCC samples after integration with Harmony. Color indicates distinct sample origins. **C.** Cluster identification of the integrated dataset, showing 12 clusters. **D.** Fractional distribution of donors across identified clusters. **E.** Fractional distribution of cancer stages (Ia or Ib) within each cluster. **F.** Fractional distribution of cell types within each cluster; tumor = cells isolated from tumor tissue; normal adjacent = cells isolated from the adjacent normal tissue; PBMC = cells isolated from peripheral blood mononuclear cells. **G.** UMAP representation highlighting different cell origins listed in panel F. **H.** Enrichment of the Tr1-specific TF motif signature using Gene Set Enrichment Analysis (GSEA). NES = normalized enrichment score. *q*-value = FDR-corrected *p*-value. Color scale = *q*-value-adjusted NES. **I.** UMAP embedding of accessible motifs for indicated transcription factors (TFs) based on chromVAR deviation scores, calculated by ArchR. **J.** UMAP embedding of accessible genes based on gene score accessibility per cell, as calculated by ArchR.

Like in the peripheral blood CD4^+^ T cells (**Fig.5H-I**), CTLA4 accessibility was lower in the tumor Tr1 cluster C6 than in blood and tumor FOXP3^+^ Tregs (**Fig.6J, Table S12**), and IL10 accessibility higher, but not specific to cluster C6. Perforin (PRF1) accessibility was notable in Th1 cluster C3 and IFN-activated cluster C4 (**Fig.6J, Table S10**). We also observed that cluster C5 had a NES > 1, but enrichment was not statistically significant (**Fig.6H**). To ensure that we correctly define C6 as a Tr1 cell cluster, we examined the marker genes of clusters C5 and C6 (**Table S12**). The accessibility of IL17RE was highest in cluster C5 (**Fig.6J**), likely indicating Th17-like cells. In contrast, cluster C6 had significantly more accessible genes encoding granzyme K (GZMK, **Fig.6G**), the TF CEBPD (detected in Tr1 cells in single cell analysis, **Fig.1J**), chemokine receptor CCR5, which is highly expressed in human gut Tr1 cells^36^, and genes found in Tr1 cell DORC or in the GREAT enhancer analyses (CXCR6, CCL3, and TF ZEB2, **Fig.3G**). Interestingly, accessibility of CXCL13 was elevated in cluster C6. CXCL13 can be expressed by Tfh and Tph cells^68^, but cluster C6 did not have high accessibility of the Tfh marker CXCR5 or the Tph TF SOX4^68^ (**Fig.S7C**). These data support the identification of cluster C6 as a Tr1 cell cluster (**Fig.6J**). Altogether, we show that a TF signature, which is unique to antigen-specific human Tr1 cells induced *in vitro,* can be used to identify putative Tr1 cells within tumor-resident CD4^+^ T cells.

## Discussion

Systematic investigation of the epigenetic landscape of human Tr1 cells and their transcriptional regulators has not been performed, despite increasing interest in Tr1 biology and clinical applications in various antigen-driven inflammatory conditions and diseases^6-15^, and ongoing clinical trials of Tr1-enriched T-allo10 cell therapy in allo-HSCT^29,32,33^. Here, we show that human *in vitro*-induced antigen-specific Tr1 cells have a unique transcriptome, oligoclonal TCR repertoire, and a distinct signature of open chromatin, TF motifs, and TF footprints in comparison to the parental CD4^+^ T cells, as well as other CD4^+^ cell populations. Integrating the epigenetic data with the Tr1 cell transcriptome enabled us to identify IRF4, BATF, and MAF as top transcriptional regulators of Tr1 cells. Using functional genomics, we confirm that IRF4, BATF and MAF play critical and non-redundant roles in Tr1 cell differentiation, phenotype, and functions. We demonstrate that the TF signature discovered in antigen-induced Tr1 cells *in vitro* can also track Tr1 cells *in vivo*, in CD4^+^ T cells isolated from peripheral blood of T-allo10 treated patients. Furthermore, this TF signature can identify Tr1 cells in CD4^+^ T cells present in solid tumor tissue.

Current lack of wide consensus on the identity of Tr1 cell-regulating TFs may result from the use of different experimental models. In the mouse, where Tr1 differentiation is IL-27-dependent, systems biology- and mechanistic studies have been performed to identify their key transcriptional regulators. Polyclonal Tr1 cell induction model pointed to Irf1, Batf, Maf, and Ahr as murine Tr1 lineage-defining TFs^20-22^, antigen-specific induction model pointed to Eomes, Tbx21, and Prdm1^23,69^, while peptide-nanoparticle induction model pointed to Bcl6, Irf4 and Prdm1^70^. In humans, Tr1 cell induction is IL-10 dependent, and mechanistic studies to validate the functional role of lineage-defining Tr1 TFs are limited. In one study, lentiviral vector mediated overexpression of EOMES in CD4^+^ T cells induced polyclonal Tr1-like cells^26^, suggesting its role in Tr1 differentiation. Another study used shRNA and small molecule inhibitors to show that IRF4 cooperates with AHR to regulate IL-10 production and suppressive function of activin-A-induced polyclonal Tr1 cells^27^. We investigated the lineage-defining TFs in human antigen-specific Tr1 cells, which are induced *in vitro* in response to alloantigens presented by allogeneic tolerogenic dendritic cells^8,29,71^. Our integrative analysis of the epigenome and transcriptome of human antigen-specific Tr1 cells showed that Tr1 cells utilized IRF4, BATF, MAF, TBX21, EOMES and PRDM1, but not BCL6 or AHR as suggested in murine nanoparticle^70^ and human activin-A^27^ induction models, respectively, likely because of the intrinsic differences in these models.

Our functional genomics studies showed that IRF4, BATF and MAF are key regulators of antigen specific Tr1 cells. While IRF4 and BATF often co-operate^60^, we show that IRF4 is necessary for Tr1 cell differentiation and CTLA-4 expression, and BATF regulates granzyme B expression in Tr1 cells. We found that BATF is also highly expressed in FOXP3^+^ Tregs, which did not express granzyme B in the condition we tested. Hence, it is possible that BATF plays a different role in FOXP3^+^ Tregs than in Tr1 cells. This finding is in line with other reports suggesting that lineage-defining TFs of one CD4^+^ T-helper (Th) subset can have various functions in other Th subsets ^72^. In addition, IRF4 and BATF have been implicated in T cell activation^73^, prevention of exhaustion^61^, and Th17 differentiation^74^. It is possible that the activation of IRF4 and BATF in developing Tr1 cells does not lead to Th17 differentiation due to concomitant activation of EOMES and TBX21, which repress Th17 phenotype^23,75,76^. IRF4 also regulates the differentiation of T-follicular helper (Tfh cells)^77^ and production of their hallmark cytokine IL-21^78^. In our hands, antigen-specific Tr1 cells express IL-21 and genes associated with Tfh cells such as ICOS, but do not have accessible the TF BCL6, master regulator of Tfh cells^79,80^. This may result from high levels of PRDM1 (encoding Blimp-1) in Tr1 cells, which can repress BCL6^79^.

We identified MAF as another key transcriptional regulator of antigen-specific human Tr1 cells. MAF is a universal regulator of IL-10 production across different CD4^+^ T cell subsets^81^. In mice, Maf cooperates with Ahr and Prdm1 (encoding Blimp-1) to achieve optimal transcription of IL-10^82^. In addition, murine Maf and Prdm1 cooperate to upregulate the expression of a co-inhibitory receptor module that includes LAG3^83^, an archetypal Tr1 cell gene. Interestingly, while we show that MAF regulates IL-10 production in human Tr1 cells, MAF KO or OE did not change the expression of LAG3. This may be because PRDM1, which is highly expressed in Tr1 cells, compensates for the lack of MAF to upregulate LAG3. Furthermore, it is conceivable that MAF can play a role in promoting human Tr1 cell differentiation that is distinct from its effect on IL-10 production. Indeed, we show that even transient OE of MAF significantly increased Tr1 cell yields after their antigen-induced differentiation.

Our study also provides a novel way to identify Tr1 cells *in vivo*. Co-expression of CD49b and LAG3 has been shown to identify Tr1 cells in human peripheral blood^65^. Studies of tissue Tr1 cells proposed that the co-expression of PD-1 and CCR5 identified human Tr1 cells in the gut^36^, and EOMES expression identified Tr1 cells that infiltrate human solid tumors^14^. These discrepancies in the phenotype of blood and tissue Tr1 cells may stem from their adaptation to the local microenvironment, without changes in the core Tr1 cell identity and function. Indeed, *ex vivo* isolated Tr1 cells from blood^65^, gut^36^, and tumor tissue^14^ all suppressed Teff cell proliferation and produced IL-10 and IFN-γ, suggesting that their key transcriptional regulators are likely similar. Accordingly, we show that Tr1 cell-specific TF signature, which we identified in *in vitro*-induced antigen-specific Tr1 cells, can be used to detect Tr1 cells *in vivo*.

TF motif-based digital cytometry showed that we can quantify Tr1 cells using their epigenetic signature in a heterogenous bulk population of total peripheral blood CD4^+^ T cells, isolated from patients treated with Tr1-enriched T-allo10 cell therapy up to 300 days post-treatment. Previously, we used TCR clonotype-based tracking to demonstrate that Tr1 cells from infused T-allo10 cell products persist long term in patients^29^, but these data did not inform us on their phenotypic stability. Herein, our data indicates that the molecular signature of Tr1 cells is stable. We also showed enrichment of the Tr1 TF signature in peripheral blood of healthy donors, indicating that the Tr1 TF signature we identified in *in vitro*-induced antigen-specific Tr1 cells is a general signature of Tr1 cells, independent from antigens and *in vitro* vs *in vivo* conditions. This is further supported by the identification of a putative Tr1 cell cluster among resident tumor CD4^+^ T cells of patients with kidney cancer^67^. As ATAC-seq profiling is becoming more accessible, the Tr1-specific TF signature could be applied to study antigen-specific Tr1 cells in tumor biology and clarify their role in solid tumors^14^ vs. hematological malignancies^15^.

Our study has some limitations. IRF4 regulates the balance between glycolysis and oxidative phosphorylation in T cells^73^. It is conceivable that IRF4 KO impairs Tr1 cell differentiation due to a metabolic impairment, which cannot sustain energy-demanding clonal expansion of Tr1 cells. However, we did not observe IRF4 activity in control Teff cells, which also clonally expand in response to alloantigens^29^. Thus, it is likely that IRF4, alone or in association with BATF, also regulates metabolism-independent facets of Tr1 cell biology. Furthermore, the TF signature of Tr1 cells has been derived from *in vitro*-induced antigen-specific Tr1 cells, which allowed us to validate the top identified TFs via functional genomics. While we can use this signature to identify cells with Tr1 phenotype *in vivo*, we have tested it in a limited set of conditions. Finally, because Tr1 cells are also cytotoxic and similar to Th1, T_EMRA_ and Tph cells, verifying the GSEA-identified Tr1 cells by careful cluster annotation is warranted. Further validation of this TF signature will occur as more ATAC-seq datasets of human CD4^+^ T cells become available.

In summary, we describe the epigenetic landscape of antigen-specific Tr1 cells, and identify their unique signature of actively used TFs. This signature contains key transcriptional regulators of Tr1 cells - TFs IRF4, BATF, and MAF - and can be used to identify Tr1 cells in epigenomics datasets *in vivo*. Thus, the data provided herein illuminates two key challenges in Tr1 cell biology: their transcriptional regulation and their identification. These data can be used to develop Tr1 cell-based therapies using TF engineering, inhibit Tr1 cell differentiation *in vivo* using targeted TF degradation approaches, and serve as a basis for mechanistic studies of Tr1 cell biology. In addition, the discovery of a Tr1-specific epigenetic signature will advance our understanding on the role of Tr1 cells in autoimmune disease, inflammatory conditions, and cancer.

## Materials and methods

### Primary cells

Human peripheral blood mononuclear cells (PBMC) were sourced from buffy coats of deidentified donors (Stanford Blood Center, Palo Alto, CA, USA) and isolated using Ficoll-Paque Plus (GE Healthcare) density gradient. CD4^+^ T cells were isolated from PBMC using Human CD4+ T Cell Isolation Kits (Miltenyi Biotec, Inc., Bergisch Gladbach, Germany). Naïve CD4+ T cells were isolated using Human Naïve CD4+ T Cell Isolation Kit II (Miltenyi Biotec). CD14^+^ monocytes were isolated using Human CD14 Microbeads (Miltenyi Biotec). Cell purity was verified by flow cytometry. CD4^+^ T cells were cultured in a T cell medium of X-VIVO-15 (Lonza, Basel, Switzerland) with 5% Human AB Serum (MilliporeSigma, Burlington, MA, USA). Cells isolated from patients and allo-HSCT donors enrolled in the T-allo10 clinical trial (ClinicalTrials.gov Identifier: NCT03198234) were collected in accordance with the Administrative Panels on Human Subjects in Medical Research, Stanford University T-allo10 eProtocol #38734, after an informed consent. Clinical study design and patient inclusion and exclusion criteria can be found at ClinicalTrials.gov. Demographics, treatment and clinical data of the patients were reported previously^32^.

### Flow cytometry

Purified primary cells, DC-10 and mDC cells, T-allo10 cells, and cells used for T cell degranulation and suppression assays were stained to assess purity, phenotype, and/or proliferation via flow cytometry. Cells were first stained for viability for 15 minutes at room temperature using 100 uL of a 1:1000 (Live/Dead InfraRed) or 1:500 (Live/Dead Aqua and Live/Dead Violet) fixable viability dye in PBS. Cells were then washed and incubated for 5 minutes at room temperature (RT) with 5 uL of Fc receptor blocking reagent (Miltenyi) in 50 uL of staining buffer composed of PBS with 0.02% sodium azide and 2% fetal bovine serum, then incubated for 15 minutes at RT using an extracellular antibody cocktail with staining buffer added to 100 uL. Cells were then washed and acquired on a flow cytometer, or fixed and stained intracellularly. For cytoplasmic antigens (e.g., cytokines, intracellular proteins like granzyme B), cells were fixed with BD Cytofix or Cytofix/Cytoperm Buffer (BD Biosciences, San Jose, CA, USA) at +4°C (min. 20 min up to overnight), washed with saponin-based BD Permeabilization Buffer, then stained with intracellular antibody cocktail diluted in 50 uL BD Permeabilization Buffer for 30 min at +4°C. After intracellular staining, cells were washed with Permeabilization Buffer, and resuspended in Staining Buffer before acquisition. For intranuclear antigens (e.g., transcription factors), cells were fixed, permeabilized and stained using Pharmingen Transcription Factor Buffer (BD Biosciences) or eBioscience FOXP3 / Transcription Factor Staining Buffer Set (Invitrogen) following manufacturer instructions. Samples were acquired on a BD FACSymphony A5 or a Beckman-Coulter CytoFLEX. Flow cytometry data was analyzed using FlowJo v10.8 (BD). Antibodies for each panel are listed in **Table S14.**

### *In vitro* Tr1 cell differentiation

The T-allo10 protocol from Chen P and Cepika AM et al.^29^ was adapted to accommodate gene expression modulation with CRISPR. Briefly, tolerogenic dendritic cells (DC-10) and mature DC (matDC) were differentiated from CD14^+^ monocytes using IL-4, GM-CSF, and IL-10 or MPLA, respectively. For knockout experiments, CD4^+^ T cells were rested for 72 hours after electroporation in T cell media supplemented with 50 U/mL rhIL-2, then collected and plated 10:1 with allogeneic DC-10s in 24-well plates at a concentration of 7 × 10^5^ to 1 × 10^6^ T cells/mL in T cell media with 10 U/mL rhIL-10 (Gibco, Thermo Fisher Scientific). For CRISPRa experiments, naive CD4^+^ T cells were rested for 4 hours after electroporation in T cell media, washed, and plated 10:1 with allogeneic DC-10s in 24-well, 48-well, or 96-well flat-bottom plates at a concentration of 7 × 10^5^ – 1 × 10^6^ T cells/mL in T cell media with 10 U/mL rhIL-10 (Gibco). For both workflows, half of the medium volume was replaced at day 5 of coculture with T cell media supplemented with rhIL-10 for a final concentration of 10 U/mL. Medium was replaced with base T cell media as necessary up until day 10 of co-culture, at which point cells were collected.

### T cell anergy

T cell anergy assay was performed as described^29^. Briefly, T-allo10 cells or control T-allo Teff cells were labeled with CellTrace carboxyfluorescein succinimidyl ester (CFSE; Thermo Fisher Scientific) for 5 to 8 minutes at room temperature, and cultured alone or stimulated with allogenic or third party matDCs at 10:1 ratio, 20:1 ratio of Human T-Activator CD3/CD28 Dynabeads (Thermo Fisher Scientific) for 3 days. After culture, the cells were collected, stained for viability and surface markers, and the percentage of CFSE^-^, dividing cells within the CD3^+^CD4^+^ population was measured by flow cytometry and anergy was calculated as ((% CFSE^-^ T-allo - % CFSE^-^ T-allo10) / % CFSE^-^ T-allo) × 100.

### Degranulation assay

To assess degranulation, 100,000 to 200,000 T-allo10 cells were cultured in a 96-well plate for 6h in total of 200 uL of the base T cell medium (see Primary Cells section) with 3 ug/mL brefeldin A (Sigma Aldrich), 2 uM monensin (BD) and 5 uL of BV421 mouse anti-human CD107a antibody (clone H4A3, BD Biosciences), with or without anti-CD3/anti-CD28 Dynabeads or U937 myeloid leukemia cell line in 10:1 cell:bead ratio. After the culture, supernatant was removed by centrifugation, cells were stained for surface and intracytoplasmic antigens using the protocol above, and acquired. Antibodies for the degranulation panel are listed in **Table S14.**

### T cell suppression

Autologous, responder CD4^+^ T cells (2.5 × 10^4^ per well), labeled with 5 μM CellTrace Violet (Thermo Fisher Scientific) for 5 to 8 min at room temperature, were co-cultured with or without equal numbers of CFSE-labeled T-allo10 cells and stimulated with 2.5 × 10^3^ allogeneic matDCs for 5 days as described^29^. Both responder and T-allo10 cells were also plated alone and with anti-CD3/anti-CD28 Dynabeads (Gibco) as proliferation controls. At day 5 of this assay, the percentage of divided CellTrace Violet (CTV)-negative responder cells was assessed by flow cytometry. Suppression was calculated as follows: % suppression = ((% divided responder cells with only matDCs added) - (% divided responder cells with T-allo10 and matDCs added)) / (% divided responder cells with only matDCs added).

### Fluorescence-activated cell sorting (FACS)

Purification of T-allo10 cell populations, ex vivo isolated CD4^+^ T cells, and ex vivo Tr1 cells was performed as described^29^. Briefly, to isolate Tr1 cells and other subsets from T-allo10 cultures, cells were collected at day 10 of culture and stained as described (see Flow Cytometry section) using PBS with 2% FBS at all steps. Samples were then filtered through a 40-mm cell filter (BD). Cells were sorted using a BD FACSAria II with a 100-um nozzle by gating through singlets, live, lymphocyte, CD3^+^CD4^+^, CD45RA^-^ (memory) and finally LAG3^+^CD49b^+^ for Tr1 cells; LAG3^-^CD49b^-^ for DN cells; and LAG3^+^CD49b^-^ for Teff cells. To isolate total CD4^+^ T cells, cells were gated through singlets, live, lymphocyte, and CD3^+^CD4^+^ gate. To isolate naïve CD4^+^ T cells, cells were gated through singlets, live, lymphocyte, CD3^+^CD4^+^ and CD45RA^+^ gate,

### Sample processing and library generation for sc-Immune Profiling (RNA- and TCR-seq)

Live singlet CD3^+^CD4^+^ T cells were purified by FACS, counted, and their cell concentration adjusted in PBS with 0.04% BSA to allow capture up to 10,000 cells per well of the Chromium Next GEM chip, which was then loaded onto a Chromium Single Cell Instrument (10x Genomics, Pleasanton, CA). RNA-seq and V(D)J libraries were prepared using the Chromium Single Cell 5′ Library, Gel Beads, & Multiplex Kit according to manufacturer’s instructions. Paired-end sequencing was conducted on an Illumina NovaSeq 6000. The Cell Ranger Single-Cell Software Suite (10X Genomics) was employed for sample demultiplexing, alignment, filtering, and UMI counting for RNA-seq libraries, using the human GRCh38 genome assembly and RefSeq gene model. For V(D)J libraries, the Cell Ranger Single-Cell Software (10X Genomics) performed sample demultiplexing, de novo assembly, alignment, annotation against germline segment V(D)J reference sequences and clonotype grouping.

scRNA-seq data were processed using Seurat (v.3.0) with specific criteria: nFeature_RNA between 500 and 5000, percent.mito < 0.25, and nCount_RNA < 50,000. Additionally, cells with multiple TCR alpha or beta chains were filtered out. The filtered data were integrated, batch-effect-corrected, clustered, and analyzed using the standard dataset integration and analysis workflow in Seurat. After merging samples into a single Seurat object, gene expression counts underwent normalization and scaling. Graph-based clustering was performed with the top 20 principal components at a resolution of 0.5. Marker genes for each cluster were identified and clusters expressing canonical marker genes from different cell types were manually annotated. Differential gene expression analysis between T cell populations was conducted using Seurat FindMarkers() function.

TCR-seq data were analyzed using scRepertoire (v.1.0.2). A TCR clonotype was defined as the combination of genes in the TCR and the nucleotide sequence of the CDR3 region (gene + nucleotide) for paired TCR alpha and beta chains. We generated an overlay of the position of clonally expanded cells using clonalOverlay(). To look at the network interaction of clonotypes shared between clusters along the single-cell dimensional reduction we used the chord diagrams from the circlize R package, using functions getCirclize() and chordDiagram(). Additionally, the count of cells by cluster assigned into specific frequency ranges was obtained with the occupiedscRepertoire() function and clonal diversity was calculated using clonalDiversity().

### Analysis of expression dynamics of the Tr1 cell cluster

We used SCANPY, a python package for large-scale differential gene expression analysis^84^ (https://github.com/scverse/scanpy), to load our 10x output as anndata object. Structuring the data within an anndata matrix allowed for downstream compatibility with other python-based implementation methods. Preprocessing, filtering, clustering, and initial visualizations were performed using SCANPY followed by RNA velocity analysis using scVelo^85^ (https://github.com/theislab/scvelo). Velocyto, a command line interface^39^ (https://github.com/velocyto-team/velocyto.py), was used to prepare the data to be analyzed in scVelo by generating a spliced/unspliced expression matrix that is necessary for estimating RNA velocities. Vector fields were plotted using scVelo’s velocity_embedding_grid function to visualize expression dynamics across all genes. To compare velocity and expression values of key Tr1 genes, the velocity function was used to plot the ratio of spliced and unspliced transcript as well as UMAP plots with colored gradient values of velocity and expression.

### Bulk RNA- and ATAC-seq sample preparation and sequencing

T-allo10 and control T-allo cells were generated from total CD4+ T cell / dendritic cell donor pairs generated from five healthy donors as previously described (^29^, **Fig.2A**). Tr1 and non-Tr1 cells were isolated from T-allo10 cell products at day 10 of culture by FACS, as live singlet CD4^+^ CD3^+^ CD45RA^-^ cells either co-expressing CD49b and LAG3 (Tr1) or CD49b^-^LAG3^-^ (non-Tr1, double-negative or DN cells). Effector T cells (Teff) were isolated from T-allo10 products as live singlet CD4^+^CD3^+^CD45RA^-^CD49b^-^LAG3^-^ cells. Along with an aliquot of previously cryopreserved CD4^+^ T cells, these populations were either lysed with RNAqueous Lysis Buffer, RNA isolated and libraries prepared as described^29^, or their native DNA was tagmented and libraries prepared following the OMNI-ATAC protocol^45^, with two modifications; transposition reaction was done with 350 rpm mixing and using the transposase from the Illumina Tagment DNA TDE1 Enzyme and Buffer Kit (Illumina, San Diego, CA, USA). Libraries were quantified using KAPA Library Quantification kit (Roche, Indianapolis, IN, USA), then sequenced using Hi-seq 4000 sequencer (Illumina).

For bulk ATAC-seq of CD4^+^ T cells, CD4^+^ T cells were isolated using FACS as live singlet CD3^+^ CD4^+^ cells from previously cryopreserved PBMC from the first cohort of patients with hematological malignancies treated with T-allo10 cell infusion a day before their unmanipulated allo-HSCT (ClinicalTrials.gov ID: NCT03198234) at day 35, 90 and 300 post-treatment (n = 3), and four healthy donors. Patient characteristics have been described previously^32^. Frequency of Tr1 cells in *ex vivo* CD4^+^ T cells was measured using flow cytometry as described^29^ for all patients and 3/4 healthy donors. Live purified CD4^+^ T cells were processed using OMNI-ATAC protocol and libraries sequenced using NextSeq 500 sequencer (Illumina).

### Bulk ATAC-seq Data Analysis

Adapters were trimmed using cutadapt, and reads were mapped to the hg38 genome using bowtie2 with a maximum fragment length of 2000bp. Filtering criteria included non-mitochondrial reads, mapq > 20, and properly paired reads. Duplicates were removed using Picard tools. Peak calling was performed using macs2 with specific parameters (--shift -75 --extsize 150 –nomodel --call-summits --nolambda -p 0.01 -B –SPMR) on Tn5 insertion sites.

ChrAccR (https://github.com/GreenleafLab/ChrAccR) was utilized for differential accessibility analysis between sorted T cell populations. Regions and fragments on chromosomes chrM, chrX, chrY and regions with a coverage of less than 1 in more than 50 % of samples were removed. A count matrix was generated as insertion counts across samples at consensus peakset regions using the ChrAccR regionAggregation() function. DESeq2 was employed for calculating differentially accessible peaks and visualized using MA plots from the ggmaplot package. Transcription factor motif enrichment was calculated using ChromVAR^49^, deviation scores were obtained with the ChrAccR getChromVarDev() function. Footprint plots were generated using aggregations of insertion counts in windows surrounding all occurrences of a motif genome-wide. ChrAccR normalizes these counts using kmer frequencies and uses all insertion sites to calculate the background distribution. Differential footprint analysis was performed with TOBIAS^50^ using the function BINDetect.

We utilized merged peak files to visualize peak accessibility in the WashU Epigenome Browser (http://epigenomegateway.wustl.edu). Differential accessibility between Tr1 and the other CD4 T cells subsets was calculated as described previously, focusing on peaks overlapping with IRF4 and MAF promoter and enhancers regions predicted by TRANSFAC 2.0.

### Bulk RNA-seq data processing and analysis

Bulk RNA-seq was processed as described^29^. Briefly, sequencing reads were mapped and aligned to the reference human genome (GENCODE v32) and quantified using Salmon (v1.10.2), then aggregated to gene level, and imported into R using tximport. Samples were filtered to remove reads with low or no expression across multiple samples. Normalization, visualization, and differential gene expression analysis was performed using DESeq2 package^86^. The integration of RNA and ATAC data was accomplished through a paired analysis of corresponding donors. This involved identifying and examining the correlation between highly expressed genes in the RNA dataset and accessible genes in the ATAC dataset using an adaptation of functions of the FigR package (v 0.1.0).

### Functional Analysis of cis-Regulatory Regions

The GREAT^56^ gene ontology tool (v.4.0.4, http://great.stanford.edu/public/html/index.php) was employed to assign gene ontology terms to cis-regulatory regions with high accessibility in the Tr1 cluster (Tr1 peaks were defined as those with FDR less than 0.01 and log2FoldChange greater than 1.5 compared with other CD4 T cell clusters). We used the following parameters: Species assembly: hg38, Association rule: Basal+extension: 5000 bp upstream, 1000 bp downstream, 1000000 bp max extension, curated regulatory domains included. We also used FigR package (v 0.1.0, https://github.com/buenrostrolab/FigR) to discern cis-regulatory interactions and delineate domains of regulatory chromatin, referred to as DORCs. For visualization we use the function dorcJPlot().

### CRISPR/Cas9 knockouts

Single-guide (sg) RNAs were purchased from Synthego (Menlo Park, CA, USA). The sgRNAs were modified to incorporate 2′-O-methyl-3′-phosphorothioate bonds at the three terminal nucleotides of the 5′ and 3′ ends^87^. For each gene, 1 to 3 sgRNAs were selected using CRISPOR^88^ and used in combination. HiFi Cas9 was purchased from Integrated DNA Technologies (Coralville, IA, USA).

When using frozen CD4^+^ T cells, cells were thawed and rested in an 37°C / 5% CO_2_ incubator in the T cell medium with 50 U/mL rhIL-2 (Peprotech, Thermo Fisher Scientific, Waltham, MA, USA), for 4-24 hours prior to electroporation, while freshly isolated CD4^+^ T cells were electroporated immediately following isolation. 37.2 pmol of Cas9 and 105 pmol of sgRNA were incubated for 30’ at room temperature to form RNPs. T cells were resuspended in P3 buffer and mixed with complexed RNPs and 10 pmol of Cas9 Electroporation Enhancer (Integrated DNA Technologies). Cells were electroporated using the Lonza 4D Nucleofector X unit, program EO-115, then plated at 1 × 10^6^ to 4 × 10^6^ cells/mL in T cell media with 50 U/mL rhIL-2 and rested for 72h before lysis for Sanger sequencing analysis or T-allo10 culture initiation. CRISPR/Cas9 KO sequences are listed in **Table S15**.

### Indel frequency analysis

60-72h post-targeting, CD4^+^ T cells were collected and gDNA was isolated with Quick Extract (Lucigen) following manufacturer instructions. The following PCR primers were used to amplify cut sites with Q5 Hot Start High-Fidelity 2X Master Mix (New England Biolabs, Ipswich, MA, USA): IRF4 5’ - AGGTGCCTTCTTCCGGGG – 3’ & 5’ - TTGCGTGGAAACGAGAACGC – 3’; BATF 5’ – GAAGTTTCCGCCCATGTGAC – 3’ & 5’ – CGGCCCACTTGAAAACTCCT – 3’; MAF 5’ – CGCCGCGCAAGCTAGAA – 3’ & 5’ – GGGTAGCCGGTCATCCAGT – 3’. PCR products were Sanger sequenced using one of the primers shown above. Resulting Sanger chromatograms were then used as input for indel analysis (Synthego Performance Analysis, ICE Analysis. 2019. v3.0. Synthego; February 2023)).

### CRISPRa

Chemically modified sgRNAs for CRISPRa were selected using CRISPick^89^ (https://portals.broadinstitute.org/gppx/crispick/public). Two guides for each gene were chosen based on GC content and proximity to TSS, then used in combination. sgRNAs, chemical modifications identical to those used for CRISPR/Cas9 knockouts, were purchased from Synthego. dCas9-VPR mRNA was obtained from Horizon Discovery (Waterbeach, UK). Cells were resuspended in P3 buffer (Lonza) and mixed with 105 pmol of sgRNA and 0.6 ug of dCas9-VPR mRNA, followed by electroporation on a Lonza 4D Nucleofector X unit with program EO-115, using a Lonza electroporation strip with no more than 1.5 × 10^6^ cells per well. Cells were plated at 1 × 10^6^ to 4 × 10^6^ cells/mL in T cell media and rested before lysis for qPCR and T-allo10 culture initiation. CRISPRa guide sequences are listed in **Table S15.**

### RNA and qPCR

24-48 hours after electroporation of CRISPRa components, cells were collected and lysed using RNeasy Micro Kit RLT buffer (Qiagen, Hilden, Germany), then passed through Chromatrap gDNA removal columns (Porvair Sciences, King’s Lynn, UK). From here, Qiagen RNeasy Micro Kit instructions were followed for RNA extraction, minus DNase treatment steps. RNA yield was quantified using a Qubit 4 fluorometer (Invitrogen, Thermo Fisher Scientific) and reverse transcription carried out using SuperScript VILO IV mastermix (Invitrogen). cDNA input for qPCR was normalized between samples based on pre-cDNA synthesis RNA concentrations. Taqman Fast Universal PCR master mix and Taqman gene expression assays (Applied Biosystems, Thermo Fisher Scientific) were used for qPCR. The assay IDs of the Taqman assays used are as follows: HS00943570_m1 for RUBCN, Hs_00193519_m1 for MAF. qPCR was performed on a QuantStudio 7 Pro (Applied Biosystems) and analyzed using Design and Analysis v2.6 software (Thermo Fisher Scientific). qPCR was run with 3 technical replicates per sample. Target transcript abundance was normalized to endogenous control (RUBCN) abundance within samples before assessing CRISPRa efficiency by comparing non-targeted and targeted conditions.

### Estimation of Tr1 cell abundances with CIBERSORTx

The CIBERSORTx^64^ algorithm was employed to generate a signature matrix of accessible motifs for isolated Tr1 cells (Create Signature Matrix module). This involved analyzing motif deviations from bulk ATAC-seq data obtained from isolated CD4^+^ T cells, Tr1 cells, and non-Tr1 (DN) cells within T-allo10 products. Subsequently, the generated signature matrix was applied to predict the composition of Tr1 cells within total CD4^+^ T cells from both healthy donors and post-allo-HSCT patients, as well as following T-allo10 infusion, using bulk ATAC data (Input Cell Fractions module).

### Sample processing and library generation for sc-multiome (ATAC- and RNA-seq)

CD4^+^ T cells were purified by FACS as for the bulk ATAC-seq. Nuclei were isolated from 40,000 cells according to the Demonstrated Protocol: Nuclei Isolation for Single Cell Multiome ATAC + Gene Expression Sequencing (10x Genomics CG000365). In summary, cells were washed twice by centrifugation with PBS with 0.04% bovine serum albumin (BSA; Miltenyi Biotec), then lysed with 45 µl chilled Lysis Buffer. After 3.5 min incubation on ice, the lysate was washed with 50 µl chilled Wash Buffer, supernatants removed, and pellets washed again without disruption by adding 45 µl of chilled Diluted Nuclei Buffer to each tube. Nuclei were then resuspended in 7 µl of Diluted Nuclei Buffer, and an 2 µl aliquot used for manual counting with hemocytometer and 0.4% Trypan Blue solution. After counting, nuclei concentration was adjusted to achieve maximum targeted nuclei recovery and promptly processed for single-cell library generation using the manufacturer’s protocol. Libraries were prepared according to the Chromium Next GEM Single Cell Multiome ATAC + Gene Expression User Guide (CG000338 Rev F). Paired-end 150-bp sequencing was done either on NexSeq 500 using the v2.5high-output kit (sample from healthy donor 10242) or using Illumina NovaSeq 6000, targeting a depth of 250 million read pairs per sample. Over 5,000 nuclei were captured per donor, and samples passed the quality control post sequencing (**Table S16**).

Demultiplexed scRNA- and scATAC-seq fastq files were processed using the Cell Ranger ARC pipeline (version 2.0.0) from 10x Genomics, generating barcoded count matrices for gene expression and ATAC data. The R package ArchR^90^ (v 1.0.1) was used for downstream analysis. In ArchR, count matrices for each sample were imported and filtered for barcodes present in both scRNA-seq and scATAC-seq datasets. ArchR quality control included filtering for nuclei with 200-50,000 RNA transcripts, <1% mitochondrial reads, <5% ribosomal reads, TSS enrichment >4, and >1,000 ATAC fragments. Subsequently, the filterDoublets() function in ArchR automatically removed doublets by identifying and eliminating the nearest neighbors of simulated doublets. For visualization and cluster identification we first identify variable peaks and perform dimensionality reduction using the ArchR addIterativeLS()I function. When the Latent Semantic Indexing (LSI) approach is not enough to correct strong batch effect differences, ArchR implements a commonly used batch effect correction tool called Harmony with the function addHarmony(). For visualization we used Uniform Manifold Approximation and Projection (UMAP) embeddings. To do cell type annotation, we use prior knowledge of cell type-specific marker genes and data-driven annotation tool CIPR^91^. Gene expression was estimated from chromatin accessibility data using ArchR gene scores. Pseudo-bulk ATAC replicates were created for each cell type, and chromatin accessibility peaks were called using MACS2. Marker peaks were identified using Wilcoxon pairwise comparisons using ArchR getMarkerFeatures. For motif analysis, ArchR also calculates motif deviations using ChromVAR. Transcription Factor motif enrichment analysis on scATAC peaks was executed using the peakAnnoEnrichment() function in ArchR. Default parameters were employed, utilizing position frequency matrices from Cis-BP. Differential analysis between clusters was conducted using the getMarkerFeatures() function within the GeneScoreMatrix (for accessibility), GeneExpressionMatrix, and MotifMatrix.

### Renal cell carcinoma scATAC data processing and analysis

The fragment files corresponding to each patient from the sc-ATAC-seq dataset published in Kourtis et al^67^ were obtained from the NCBI Gene Expression Omnibus (GEO) repository under accession number GSE181062 and analyzed in the same manner as above. ArchR was used for the analysis of total CD45^+^ cells from all patients. Potential doublets were identified and subsequently removed, followed by batch correction utilizing the Harmony. Following dimensionality reduction and clustering, the cell clusters corresponding to CD20 (MS4A1)^+^ B cells, CD14^+^ monocytes, and NCAM1^+^/ FCGR3A^+^ natural killer (NK) cells were excluded from downstream analysis, focusing exclusively on T cells. Specifically, T cell clusters were identified based on the expression of CD3D and CD3E markers. Subsequently, the CD8^+^ T cell subset was selectively removed, retaining only the CD4^+^ T cell population for further analysis.

### Tr1 signature and enrichment

Gene Set Enrichment Analysis (GSEA)^46^ (version 4.2.3) was employed to compute functional enrichment scores for Tr1 cells in each patient’s dataset (including healthy donors and CD45^+^ cells from clear-cell renal cell carcinoma (ccRCC) samples^67^). This calculation method was established using the identified TF signatures specific to Tr1 cells. These signatures were determined by overlapping Tr1-accessible peaks, footprints, and differential TF motif deviations identified by chromVAR. Additionally, we incorporated the TFs contained in the Tr1 cell-regulatory domains identified using FigR^58^. The average motif deviations per cluster were exported and employed in GSEA enrichment analysis. Specifically, a list of 23 distinct Tr1 motifs served as the gene set for this analysis (**Table S10**). Each cluster was treated as a unique phenotype, and its enrichment score was determined by comparing it to the remaining clusters with phenotype permutation (1000 permutations). When the resulting FDR q-value equaled zero, it was reported as <0.001, following the convention established in GSEA that a p-value of zero (0.0) signifies an actual p-value smaller than 1 divided by the number of permutations performed. To validate the specificity of this approach and motif signature, we employed a similar methodology to assess ‘Tr1 enrichment’ by re-analyzing B cells and monocytes sourced from the same clear-cell renal cell carcinoma dataset^67^.

### Statistical Analysis

Sample sizes were not predetermined using statistical methods but were comparable to those in previous publications^29^. Except in cases of technical failure, no data were excluded from the analyses, and the experiments were not randomized. Investigators were not blinded during experiments or outcome assessment, and data collection and analysis were not performed blind to experimental conditions. Statistical analysis was carried out in GraphPad Prism 9 and 10 (GraphPad Software, Inc, Boston, MA, USA) or R v.4.0.3. Data analyzed using GraphPad Prism used non-parametric tests that do not assume equal variances between groups: Mann-Whitney or Wilcoxon test for groups of two (unpaired or paired samples, respectively), and Kruskal-Wallis or Friedman analysis of variance (ANOVA) with Dunn’s post hoc test for > 2 groups (independent or dependent samples, respectively), and Spearman rank r for correlation, with testing level (α) of 0.05. Tests were two-sided unless otherwise indicated. As applicable, center bars and whiskers represent the median with interquartile range. The number of biological replicates (i.e., donors), *n*, is indicated in figure captions for each experiment. The R packages used in the analysis of sequencing data have embedded statistical algorithms that include multiple testing correction for *p* values. False discovery rate (FDR) of 0.05 was applied. All measurements displayed were taken from distinct samples, i.e. biological replicates.

## Acknowledgements

We thank Dr. Rhonda Perriman for the critical review of the manuscript, Dr. Gretchen Ehrenkaufer from Stanford SPARK and Dr. Šimon Borna and for helpful discussions, and the staff from Stanford Genomics Facility and Chan Zuckerberg Biohub Community Access Program for library preparation assistance and sequencing. This study was funded by the National Blood Foundation Early Career Research Grant (to A.M.C), American Society for Gene+Cell Therapy Career Development Award (to A.M.C), SPARK Translational Research Grant (to A.M.C and M.P-D.; supported by the Stanford Maternal and Child Health Research Institute, and The Weston Havens Foundation), Stanford Translational Research and Applied Medicine Pilot Grant (to A.M.C), Chan Zuckerberg Biohub Community Access Initiative (to A.M.C), Stanford University School of Medicine (to M.G.R), and NIH RM1-HG007735 (to H.Y.C., W.J.G.). H.Y.C. is an Investigator of the Howard Hughes Medical Institute. A portion of the sequencing data was generated with instrumentation purchased with NIH funds (S10OD025212 and 1S10OD021763).

## Author contributions

A.M.C. designed the study, obtained funding, performed experiments, analyzed and interpreted data, wrote the manuscript, and supervised the study; L.A. and C.W. designed experiments, analyzed and interpreted data, and wrote the manuscript; M.N., M.M.M., B.C.T., P.P.C., R.A.F., and F.M. performed experiments and/or analyzed data; M.P-D. designed experiments, obtained funding, and interpreted data; F.M. and J.N. interpreted data and reviewed the manuscript; R.B., M.P. and W.J.G. interpreted data and helped obtain funding; H.Y.C. interpreted data, reviewed the manuscript, and obtained funding; M.G.R. designed the study, interpreted data, reviewed the manuscript, obtained funding, and supervised the study.

## Data availability

The sequencing data is deposited to the Gene Expression Omnibus (GEO) under the accession number GSEXXXXXX.

## Supplementary Figure Legends

**Supplementary Figure 1.**
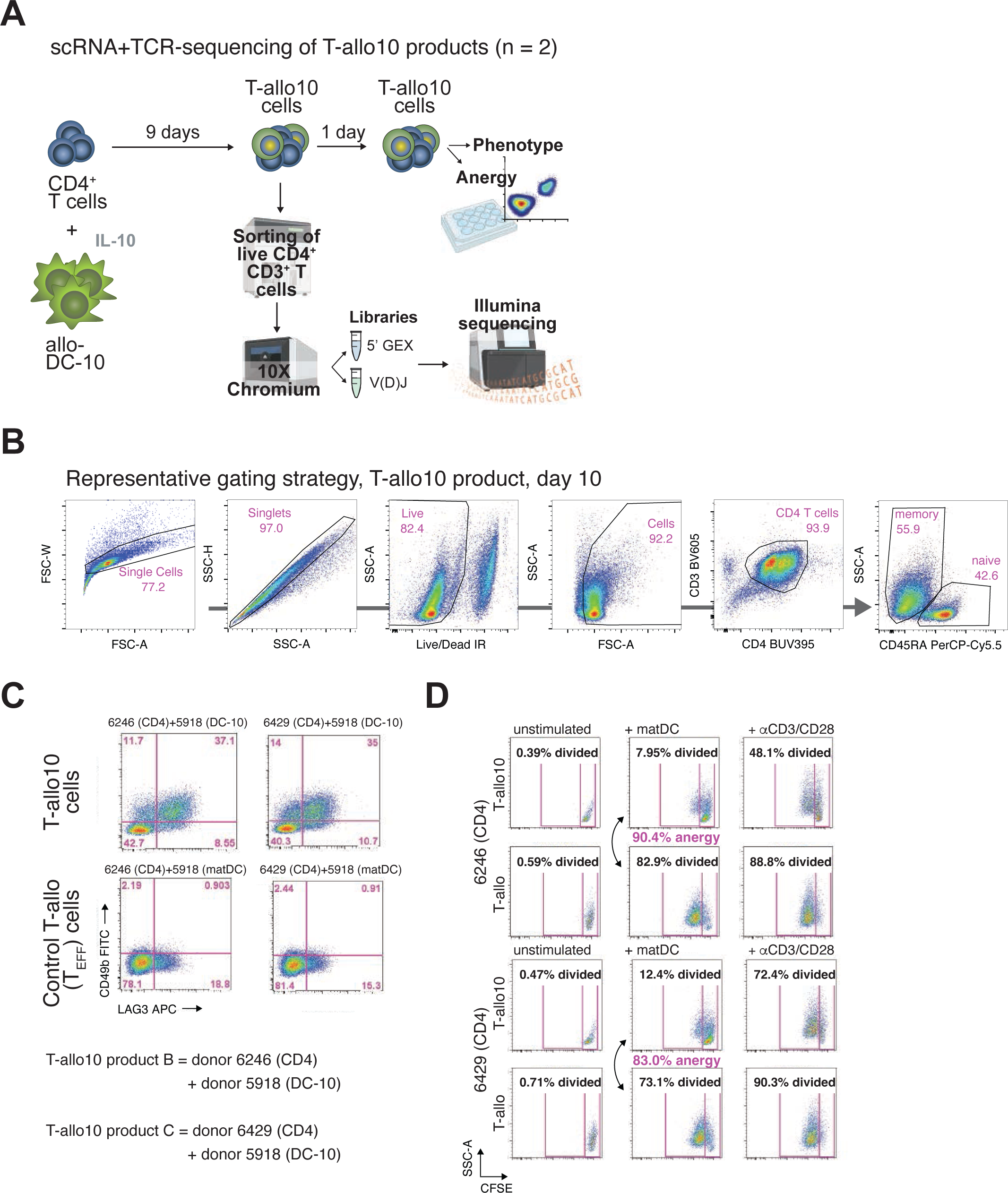
**A.** Schematic representation of 10x Genomics single-cell (sc) Immune Profiling (sc- RNA+TCR-seq) of T-allo 10 products. Aliquot of T-allo10 cells was analyzed by sc-Immune Profiling on day 9 of T-allo10 culture, while the rest was cultured by day 10 and assessed for phenotype and anergy. **B.** Representative gating strategy of T-allo10 products. **C.** Percentage of Tr1 cells in two T-allo products analyzed by flow cytometry, defined as CD49b^+^LAG3^+^ memory (CD45RA^-^) CD4^+^ CD3^+^ live singlet T cells; flow cytometry. **D.** Alloantigen–specific anergy of two T-allo10 products analyzed by flow cytometry, measured as lack of proliferation of carboxyfluorescein diacetate, succinimidyl ester (CFSE)-labeled T-allo10 cells after 3-day re-stimulation *in vitro* with allogeneic mature dendritic cells (matDC), in comparison to proliferation of control effector cells (T-allo) generated from the same parental CD4^+^ T cells but in co-culture with matDC. Anergy was calculated as described^29^. Proliferation was measured by decrease in CFSE fluorescence. Polyclonal stimulation of T-allo10 and T-allo cells with plate-bound anti-CD3 monoclonal antibody (mAb; clone OKT3, 10 μg/mL) and soluble anti-CD28 mAb (1 μg/mL) was used as a positive control for proliferation.

**Supplementary Figure 2.**
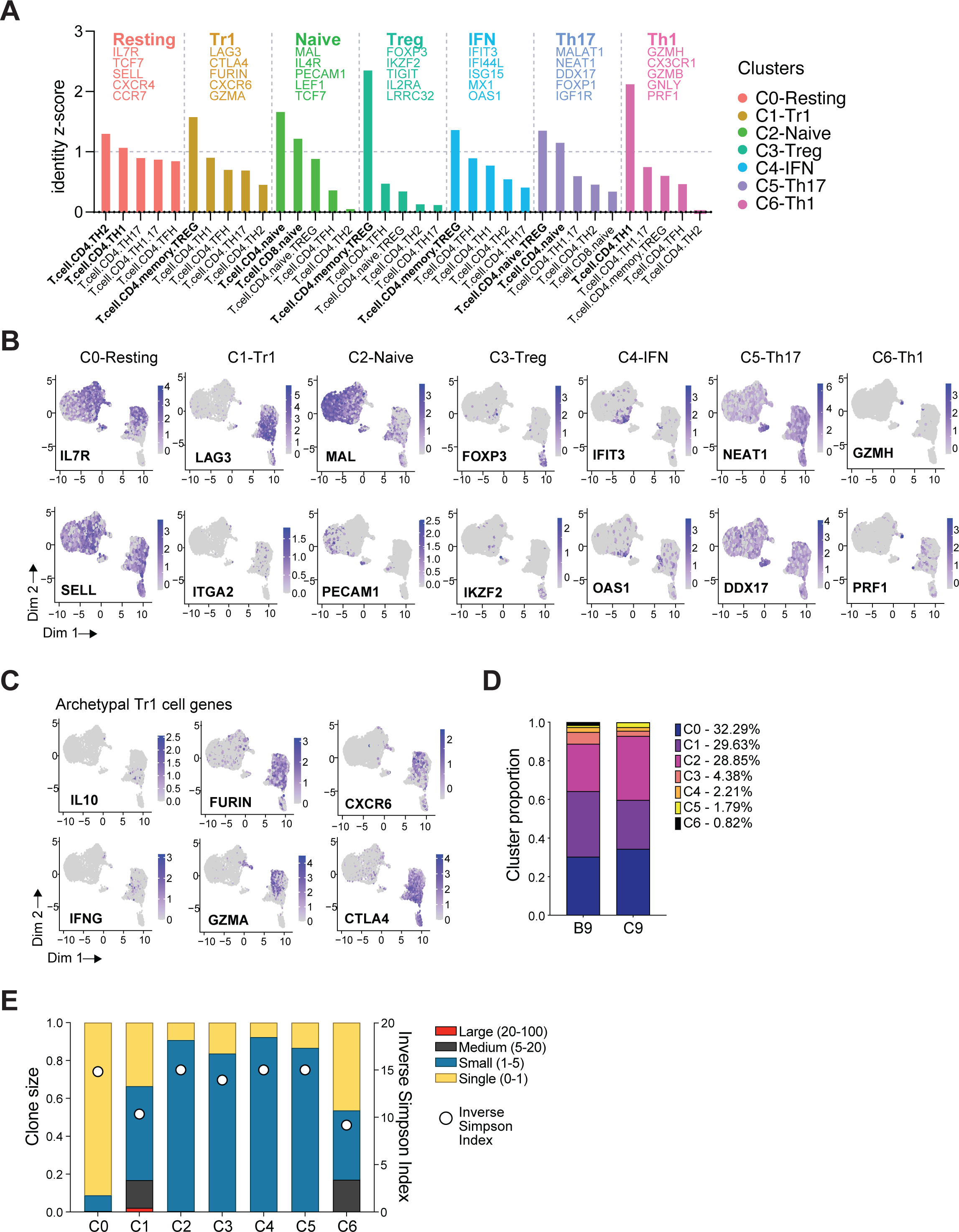
**A.** Clustering label annotation as predicted by: a) CIPR annotation tool (graph) and b) expression level of canonical markers of T cell subsets. **B.** Representative UMAP visualization of gene expression from each cluster’s defining markers. **C.** UMAP representation of gene expression from canonical Tr1 marker genes. **D.** Distribution of cluster fractions for each donor, showing the average of 2 donors. **E.** Clone size distribution of clusters and clonal diversity calculated by inverse Simpson index.

**Supplementary Figure 3.**
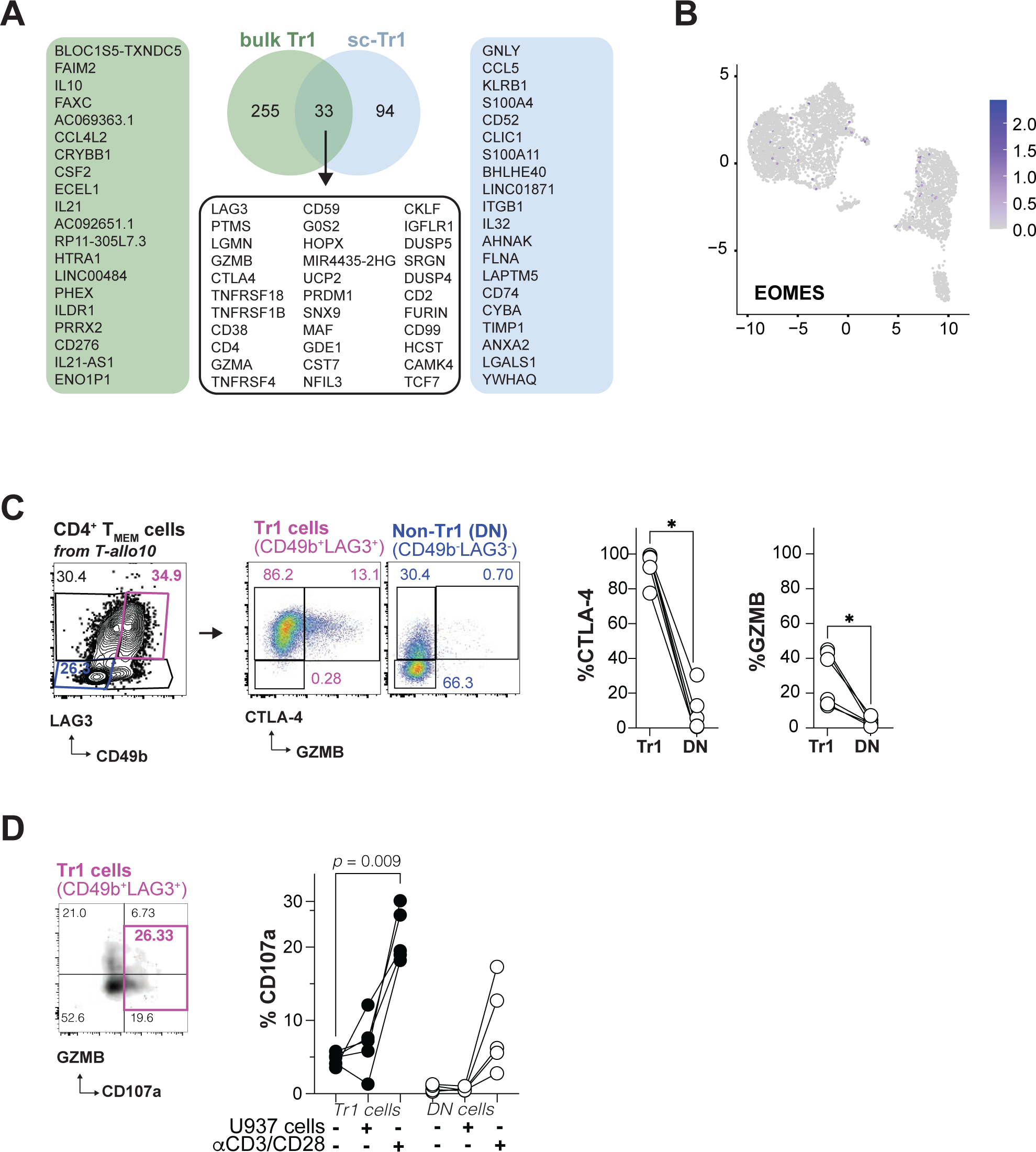
**A.** Venn diagram illustrating the overlap of upregulated genes in the T-allo10 Tr1 cell population between bulk RNA-seq (from Chen P and Cepika AM et al^29^) and single-cell RNA-seq dataset. For genes unique to bulk- and sc-datasets, top 20 upregulated genes are shown. **B.** UMAP visualization of EOMES gene expression in sc-Immune profiling of two T-allo10 products. **C.** Left: representative flow cytometry plots depicting the analysis of intracellular CTLA-4 and granzyme B (GZMB) expression in CD49b^+^LAG3^+^ Tr1 and CD49b^-^LAG3^-^ non-Tr1 (DN) memory CD4^+^ T cells from T-allo10 products. Right: cumulative data of intracellular CTLA-4 and GZMB expression, n = 6. \**p* < 0.05, Wilcoxon test. **D.** Left: representative flow cytometry plot showing the frequency of Tr1 cells expressing intracellular granzyme B (GZMB) and surface CD107a, which indicates degranulating cells, after a 6h-stimulation with anti-CD3/anti-CD28 Dynabeads. Right: cumulative data of total CD107a expression on Tr1 or non-Tr1 (DN) cells after 6h-culture of T-allo10 products (n = 5) in cell culture media alone or in the presence of anti-CD3/CD28 Dynabeads or myeloid tumor cell line U937 cells in 10:1 ratio. Friedman ANOVA with Dunn’s post hoc test was used to compare Tr1 cells or DN cells across conditions, while Wilcoxon test was used to compare Tr1 vs DN cells in each condition. Statistically significant adjusted p-values are indicated on the graph.

**Supplementary Figure 4.**
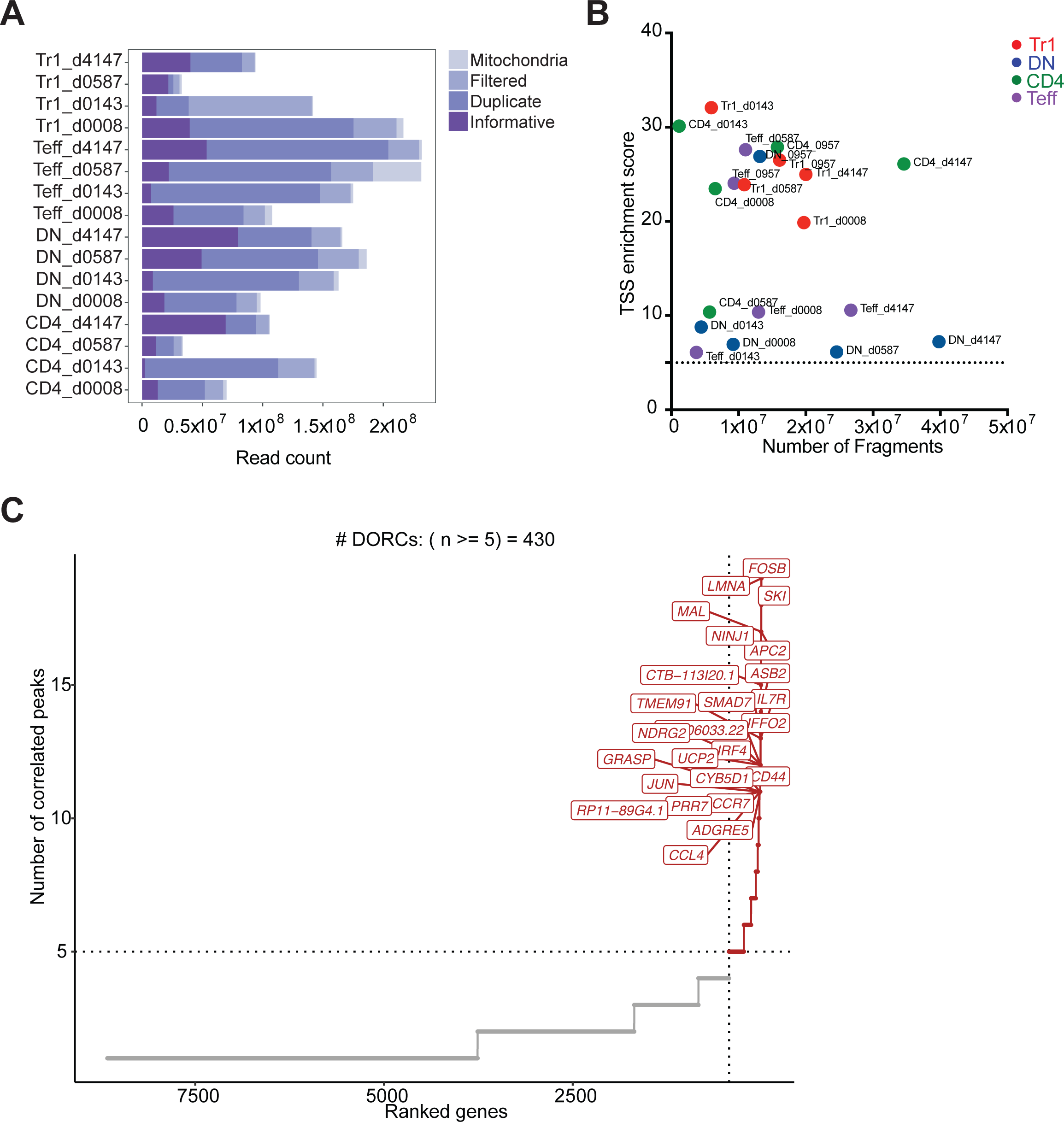
**A.** Sequence quality and unique alignment sequence distribution of the ATAC-seq dataset. **B.** ATAC-seq data quality control filters of four CD4^+^ T cell populations from five donors. Shown are the numbers of unique ATAC-seq nuclear fragments and transcription start site (TSS) enrichment in each sample. Dashed lines represent the filters for high-quality data (1 × 10^6^ unique nuclear fragments and TSS score ≥ 5). **C.** Visualization of ranked genes based on the number of significant gene-peak correlations (per gene), used to define domains of regulatory chromatin (DORC).

**Supplementary Figure 5.**
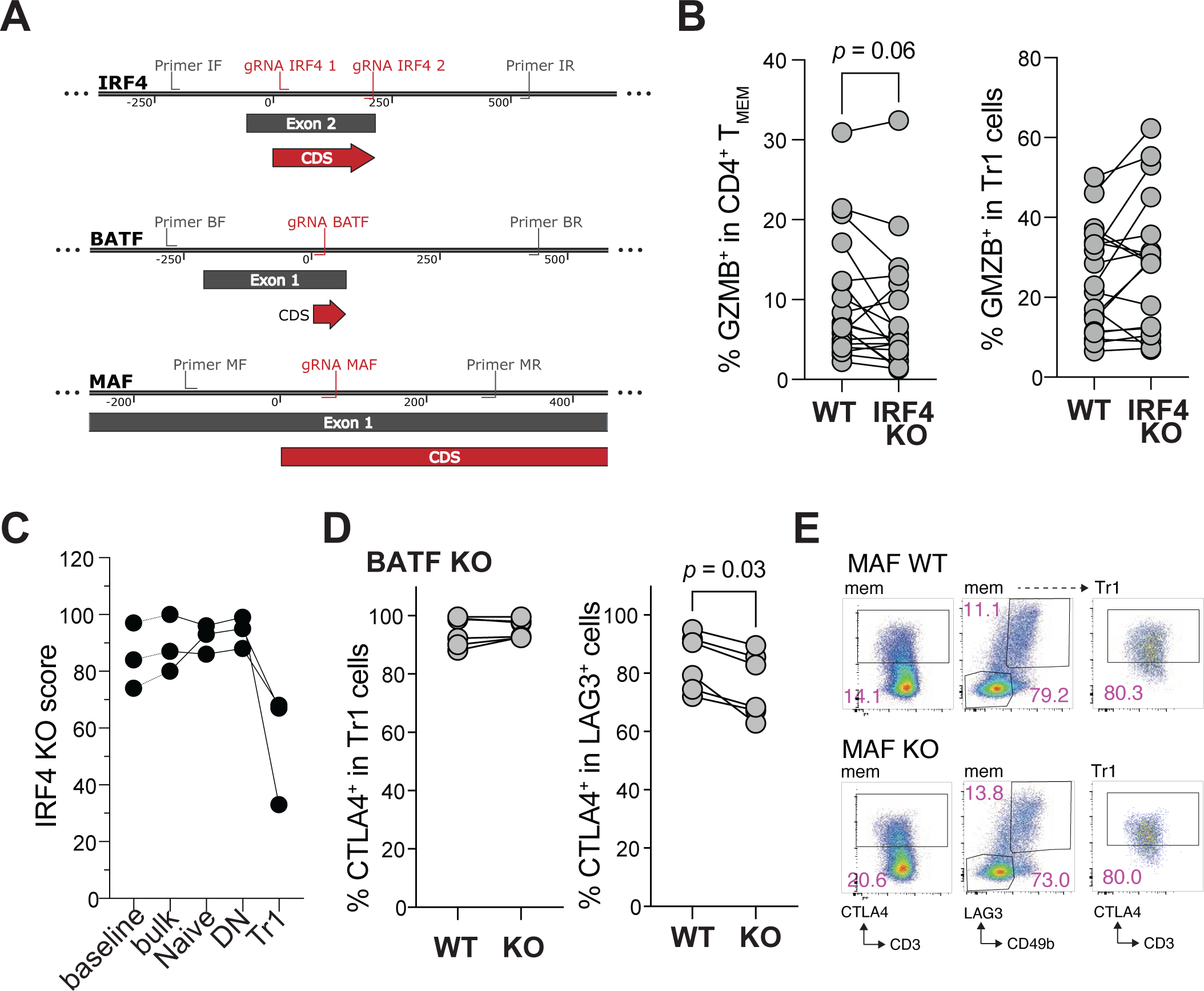
**A.** CRISPR/Cas9 single guide RNA targeting strategy for IRF4, BATF, and MAF, indicating guide RNA targets and primers used for PCR. CDS = coding sequence. **B.** Expression of granzyme B (GZMB) in memory CD4^+^ T cells (Tmem, left) and Tr1 cells (right) in wild-type (WT) and IRF4 knock-out (KO) T-allo10 cells, flow cytometry. *p* = ns, n = 25, Wilcoxon test. **C.** IRF4 KO scores, measured by Synthego ICE (Inference of CRISPR Edits) analysis of Sanger sequencing data, of parental CD4^+^ T cells after 72h rest post-eding (baseline), total T-allo10 cells after 10 day differentiation *in vitro* (bulk), FACS-sorted live singlet CD45RA^+^CD4^+^CD3^+^ T cells from T-allo10 cells (naive), FACS-sorted live singlet CD49b^-^LAG3^-^ CD45RA^-^CD4^+^CD3^+^ T cells from T-allo10 cells (DN), and FACS-sorted live singlet CD49b^+^LAG3^+^ CD45RA^-^CD4^+^CD3^+^ T cells from T-allo10 cells (Tr1). **D.** Expression of CTLA-4 in LAG3^+^CD49b^+^ Tr1 cells (Tr1, left) and in LAG3^+^CD49b^-^ memory CD4^+^ T cells (LAG3^+^, right) in wild-type (WT) and BATF knock-out (KO) T-allo10 cells, flow cytometry. n = 6, Wilcoxon test. **E**. Representative flow cytometry plots depicting frequency of CTLA-4^+^ cells and Tr1 cells in memory (mem) CD4^+^ T cells (left and middle panels, respectively), and CTLA-4 expression within Tr1 cells (right panel), in WT and MAF KO T-allo10 cells.

**Supplementary Figure 6.**
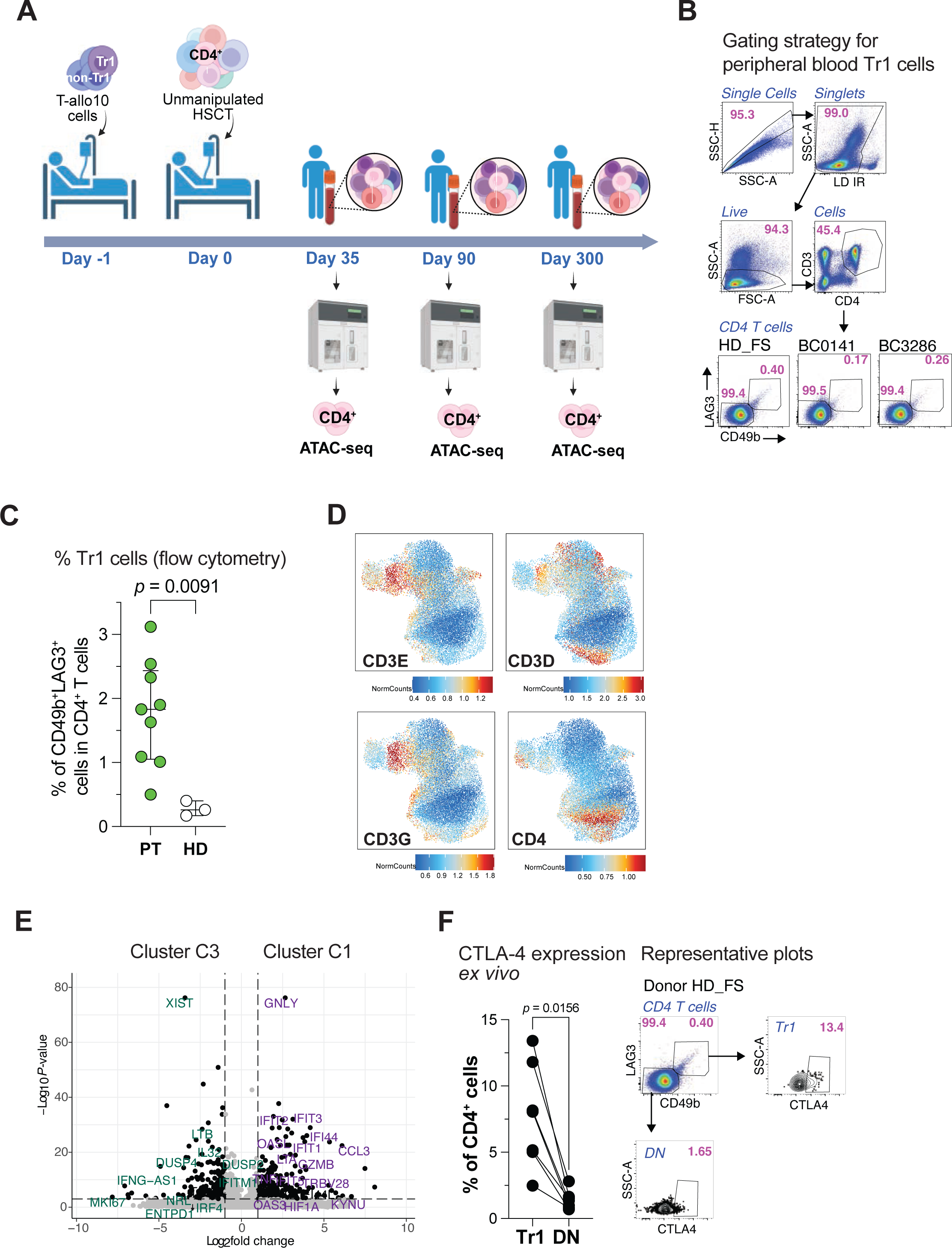
**A.** Schematic representation indicating T-allo10 adoptive cell therapy in recipients of allogeneic hematopoietic stem cell transplantation (allo-HSCT; ClinicalTrial.gov ID: NCT03198234) and time points when CD4^+^ T cells were isolated by FACS from cryopreserved patient peripheral blood mononuclear cells (PBMC) and analyzed by ATAC-seq. **B.** Top: Representative flow cytometry gating strategy to identify CD49b^+^LAG3^+^ Tr1 cells in CD4^+^ T cells within *ex vivo* PBMC. Bottom: frequency of Tr1 and CD49b^-^LAG3^-^ non-Tr1 (DN) cell populations in CD4^+^ T cells of healthy donors used for CIBERSORTx analysis. **C.** Percentage of Tr1 cells within total CD4^+^ T cells, assessed by flow cytometry in both patient (PT, n = 9) and healthy donor (HD, n = 3) samples that were also analyzed by ATAC-seq. Wilcoxon test. **D.** UMAP visualization of nuclear RNA-derived gene expression of T cell marker genes CD3 (encoded by CD3E, CD3D, CD3G) and CD4 in purified live CD4^+^ CD3^+^ T cells that were analyzed by sc-multiome and ArchR. **E.** Volcano plot illustrating differential gene expression between clusters C3 and C1 of the sc-multiome dataset. Genes highlighted in purple represent those characteristic of cluster C1, while those in green represent cluster C3. Dotted lines indicate filtering criteria with a false discovery rate (FDR)- corrected *p*-value < 0.001 and a log2 fold change > 1. **F.** Expression of intracellular CTLA-4 (n = 6) in *ex vivo* Tr1 and non-Tr1 (DN) cells within total CD4^+^ T cells from unstimulated PBMC. Left: cumulative data, right: representative flow cytometry gating; Wilcoxon test.

**Supplementary Figure 7.**
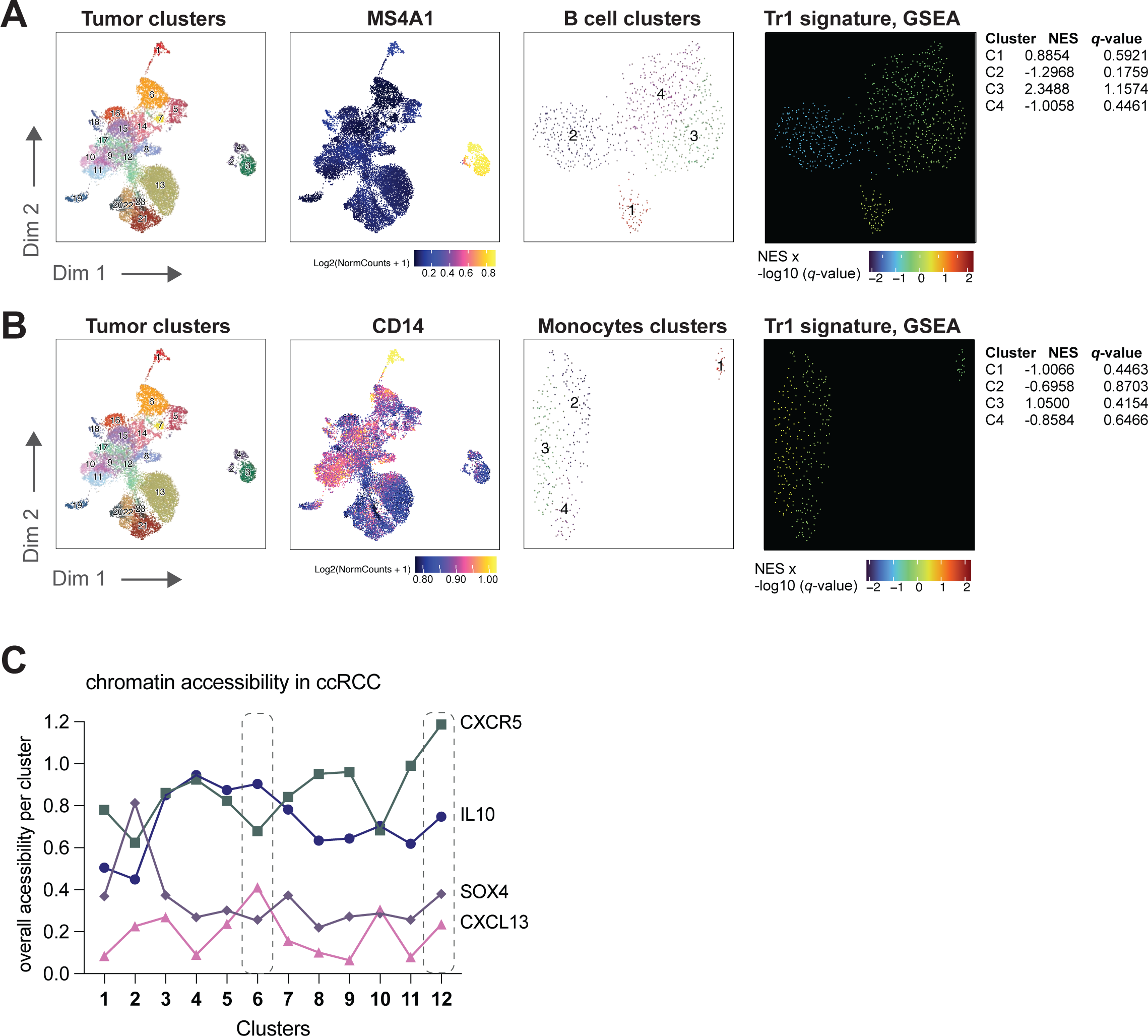
Tr1 TF signature validation in B cells (**A**) and monocytes (**B**). Analysis encompasses initial clustering of CD45+ cells analyzed by sc-ATAC-seq (from the publicly available clear cell renal cell carcinoma (ccRCC) dataset^67^) and identification of B cells (MS4A1+) and monocytes (CD14+), followed by subsequent clustering of B cells and monocytes and examination of Tr1 signature enrichment using Gene Set Enrichment Analysis (GSEA). MS4A1 gene encodes CD20.

**Table S1.**
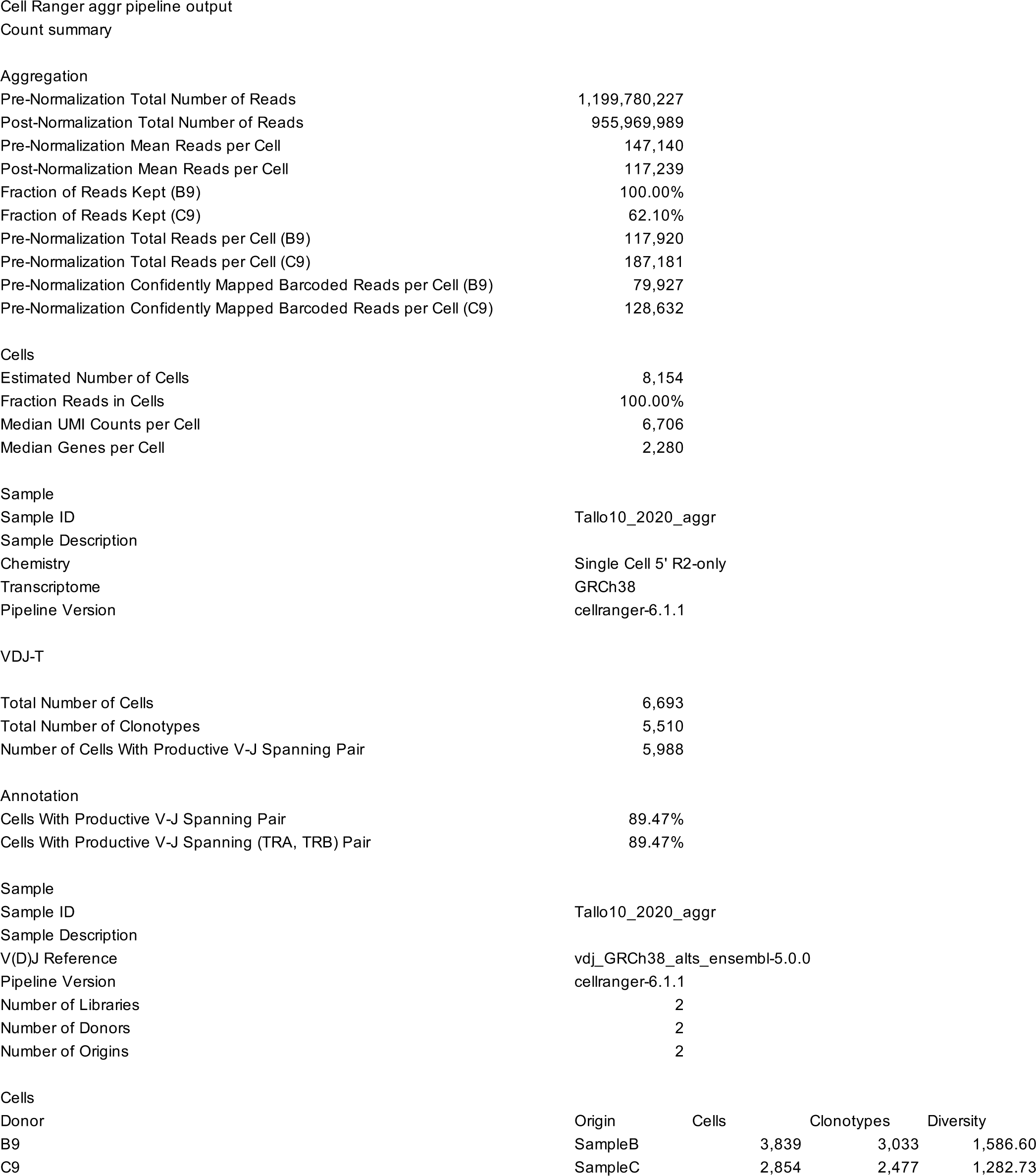
Quality control metrics of 10X single-cell Immune Profiling (RNA+TCR-seq) Related to Figure 1

|  |  |
| --- | --- |
| Pre-Normalization Total Number of Reads | 1,199,780,227 |
| Post-Normalization Total Number of Reads | 955,969,989 |
| Pre-Normalization Mean Reads per Cell | 147,140 |
| Post-Normalization Mean Reads per Cell | 117,239 |
| Fraction of Reads Kept (B9) | 100.00% |
| Fraction of Reads Kept (C9) | 62.10% |
| Pre-Normalization Total Reads per Cell (B9) | 117,920 |
| Pre-Normalization Total Reads per Cell (C9) | 187,181 |
| Pre-Normalization Confidently Mapped Barcoded Reads per Cell (B9) | 79,927 |
| Pre-Normalization Confidently Mapped Barcoded Reads per Cell (C9) | 128,632 |
| Cells |  |
| Estimated Number of Cells | 8,154 |
| Fraction Reads in Cells | 100.00% |
| Median UMI Counts per Cell | 6,706 |
| Median Genes per Cell | 2,280 |
| Sample |  |
| Sample ID | Tallo10_2020_aggr |
| Sample Description |  |
| Chemistry | Single Cell 5' R2-only |
| Transcriptome | GRCh38 |
| Pipeline Version | cellranger-6.1.1 |
| VDJ-T |  |
| Total Number of Cells | 6,693 |
| Total Number of Clonotypes | 5,510 |
| Number of Cells With Productive V-J Spanning Pair | 5,988 |
| Annotation |  |
| Cells With Productive V-J Spanning Pair | 89.47% |
| Cells With Productive V-J Spanning (TRA, TRB) Pair | 89.47% |
| Sample |  |
| Sample ID | Tallo10_2020_aggr |
| Sample Description |  |
| V(D)J Reference | vdj_GRCh38_alts_ensembl-5.0.0 |
| Pipeline Version | cellranger-6.1.1 |
| Number of Libraries | 2 |
| Number of Donors | 2 |
| Number of Origins | 2 |

| Donor | Origin | Cells | Clonotypes | Diversity |
| --- | --- | --- | --- | --- |
| B9 | SampleB | 3,839 | 3,033 | 1,586.60 |
| C9 | SampleC | 2,854 | 2,477 | 1,282.73 |

**Table S2.**
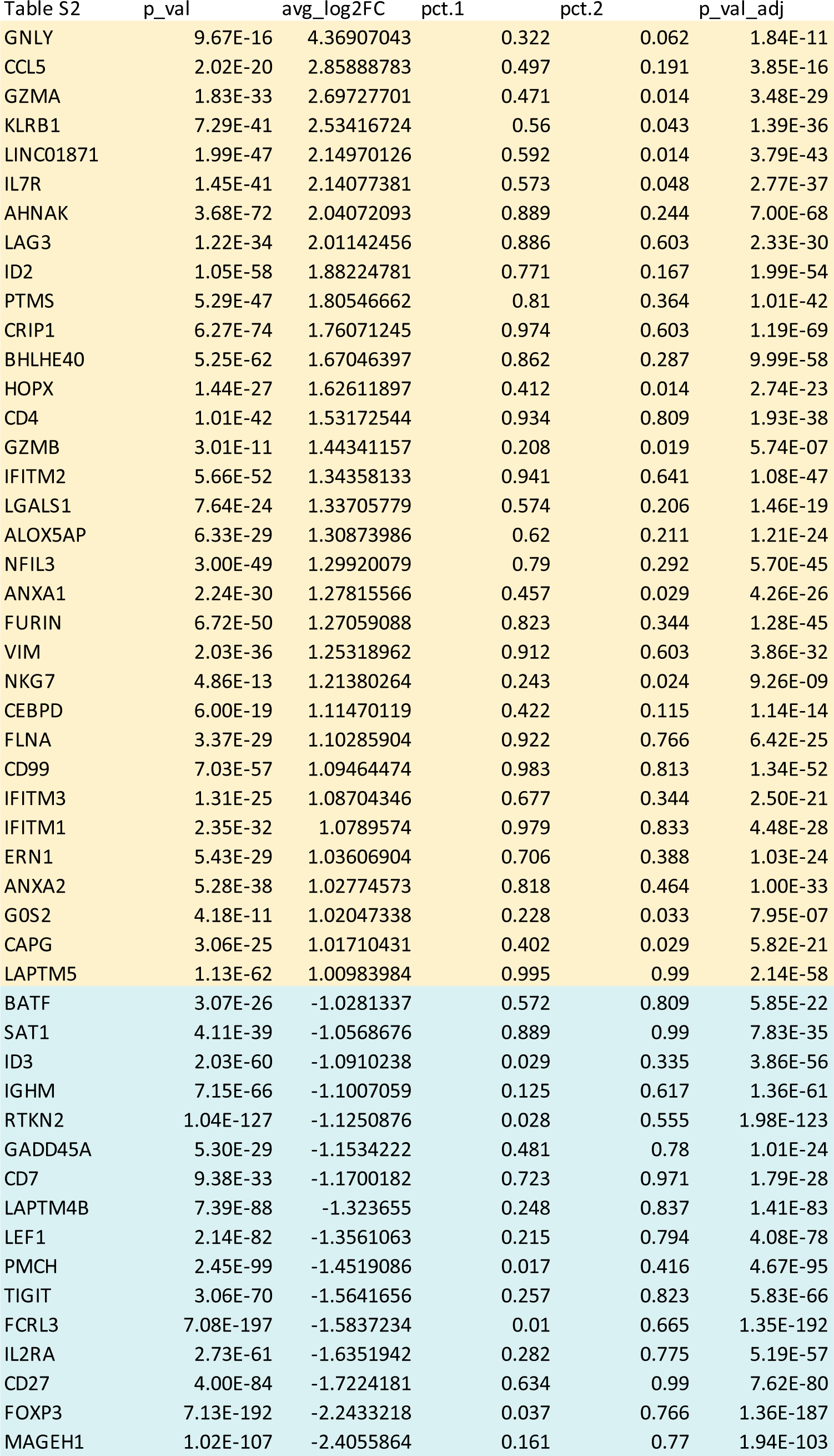
scRNA and TCR-seq, Tr1 vs FOXP3 Treg comparison. Related to Figure 1 yellow = upregulated in Tr1; turquoise = upregulated in FOXP3 Treg

**Table S3.**
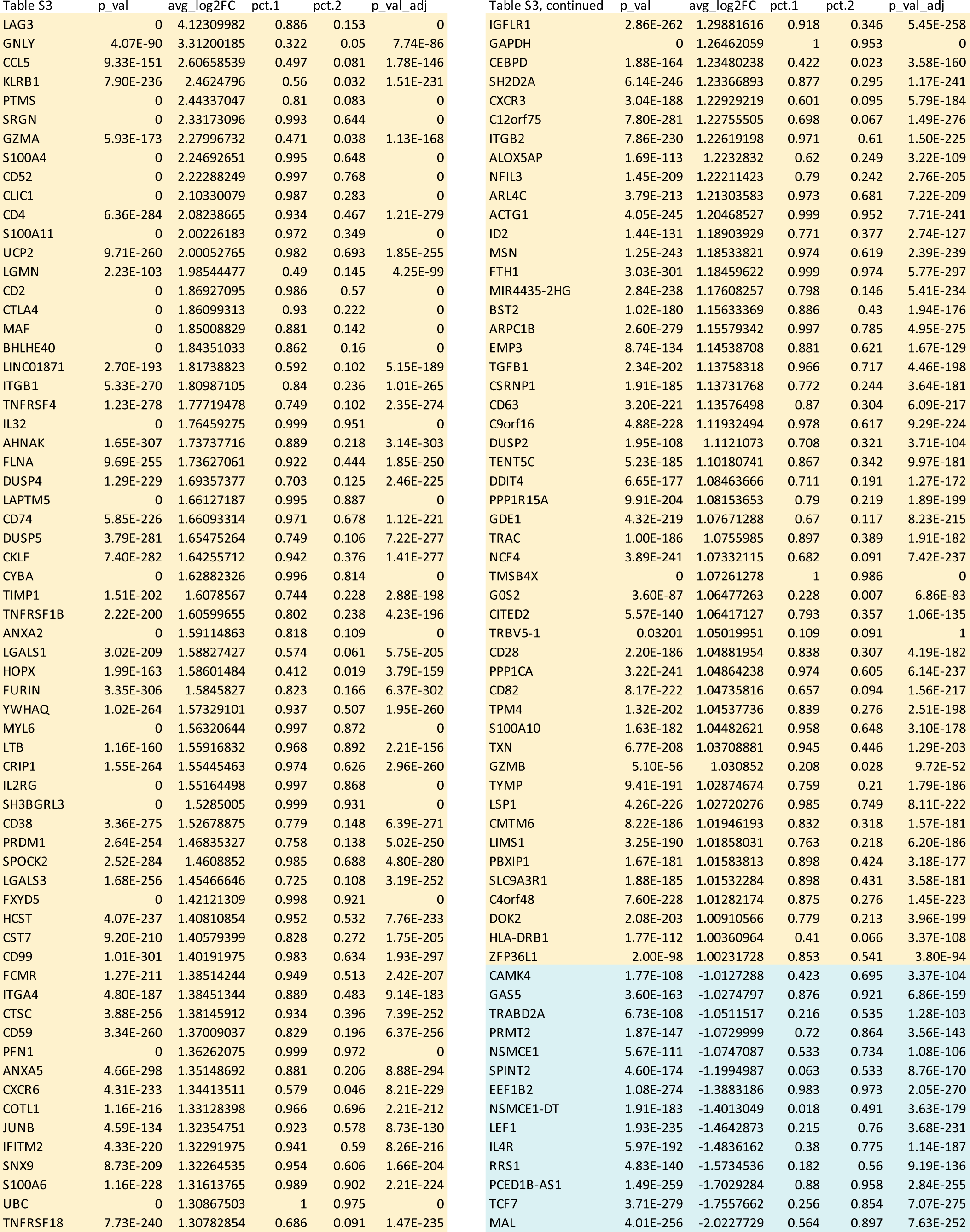
scRNA and TCR-seq, Tr1 vs all other clusters comparison. Related to Figure 1 yellow = upregulated in Tr1; turquoise = upregulated in all other clusters

**Table S4.**
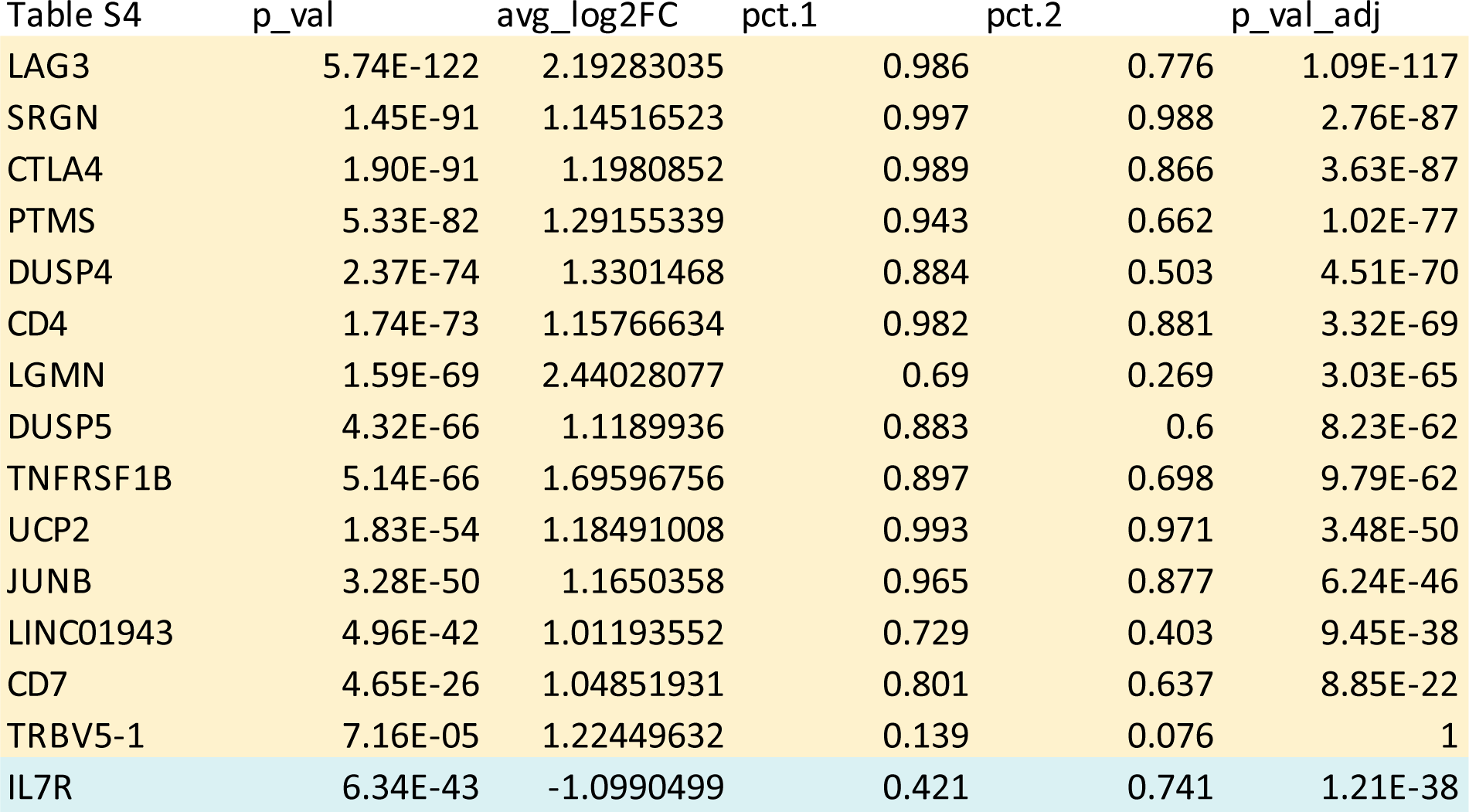
scRNA and TCR-seq, Tr1-B vs Tr1-A cluster comparison. Related to Figure 1 yellow = upregulated in Tr1-B; turquoise = upregulated in Tr1-A cluster

**Table S5.**
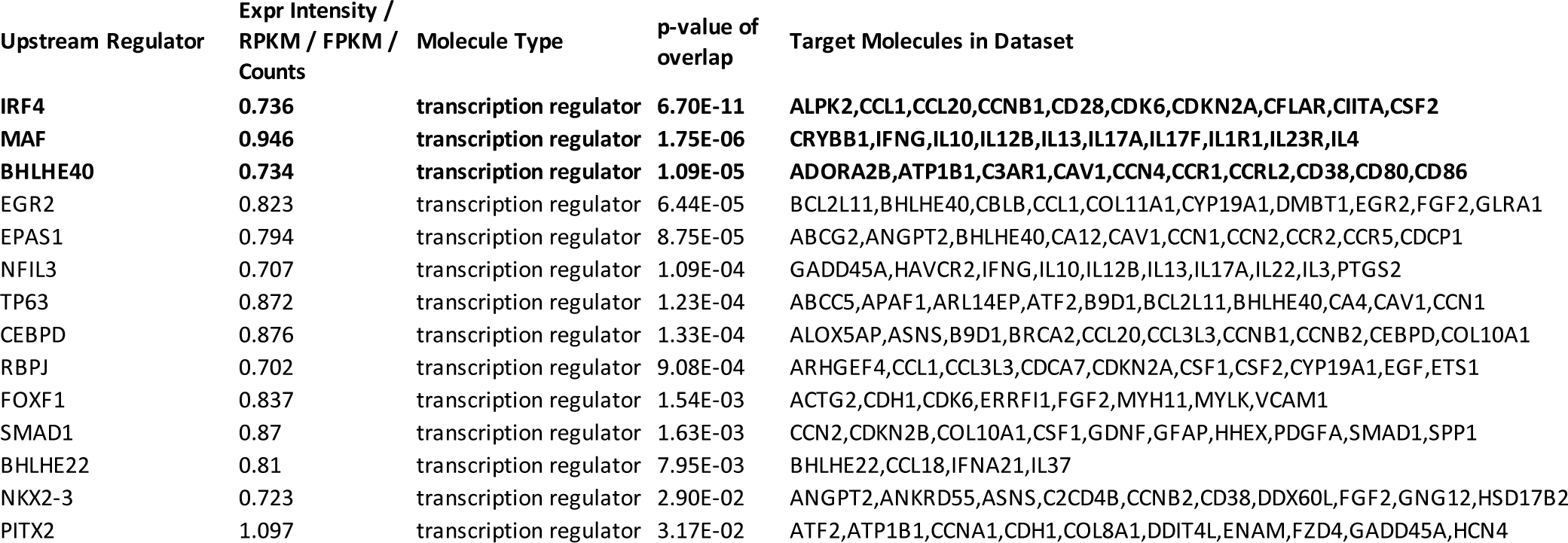
Ingenuity Pathway Analysis (IPA) upstream regulator analysis. Related to Figure 2 The list of upstream regulators was filtered by Molecule Type = transcription regulator with accessibility in Tr1 cells vs all others > 0.7 and p-value of overlap < 0.05.

**Table S6.**
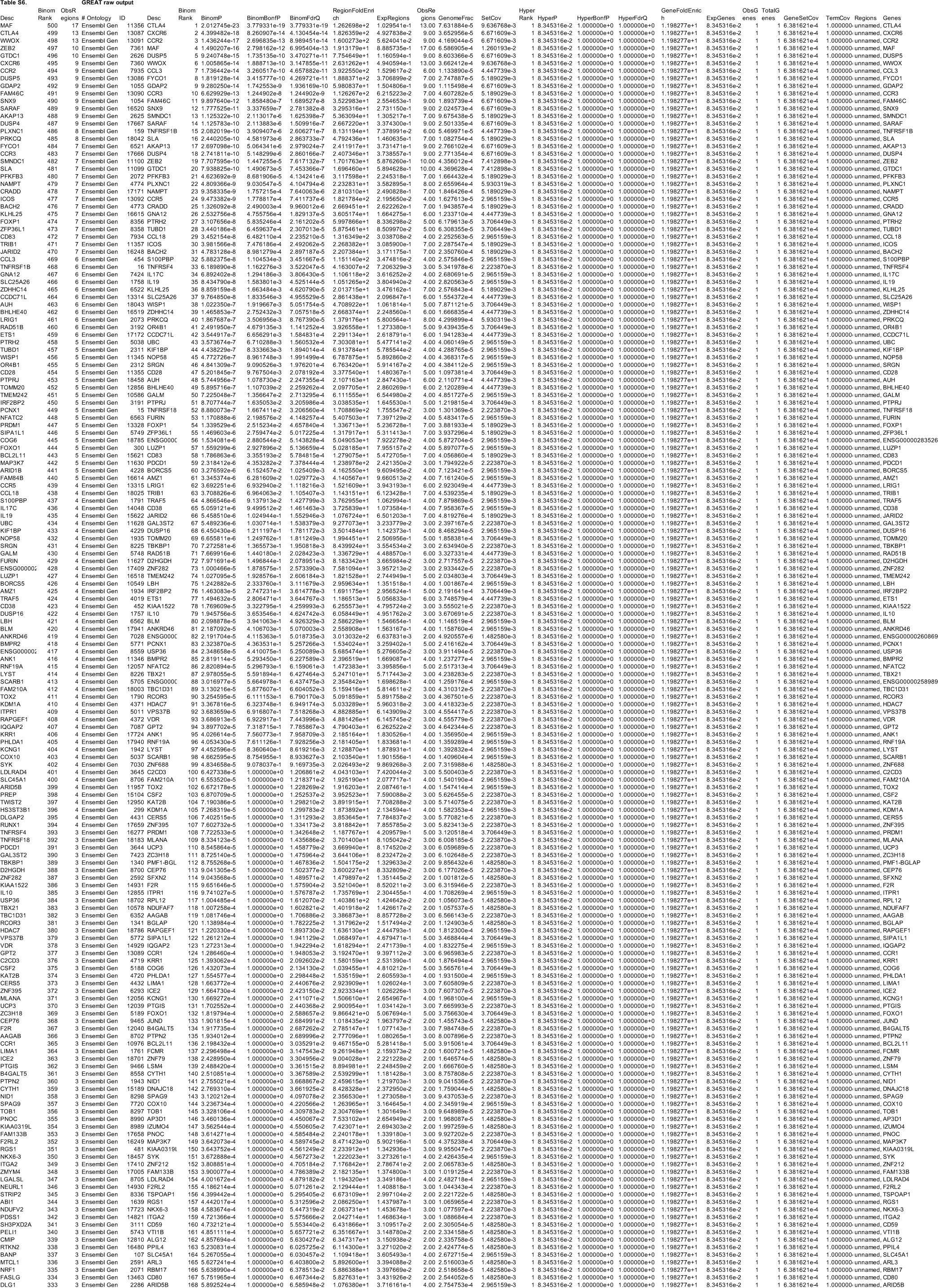
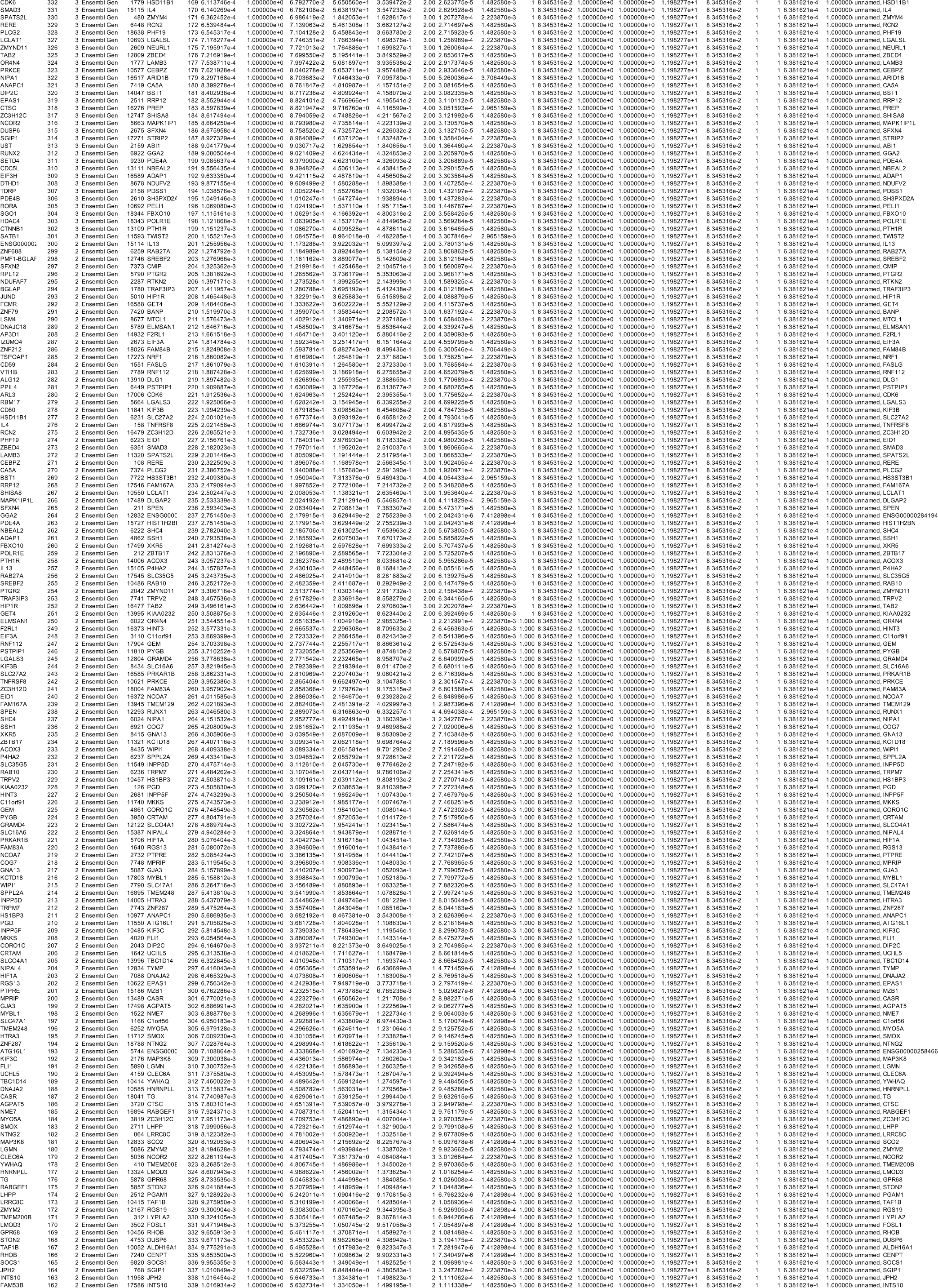
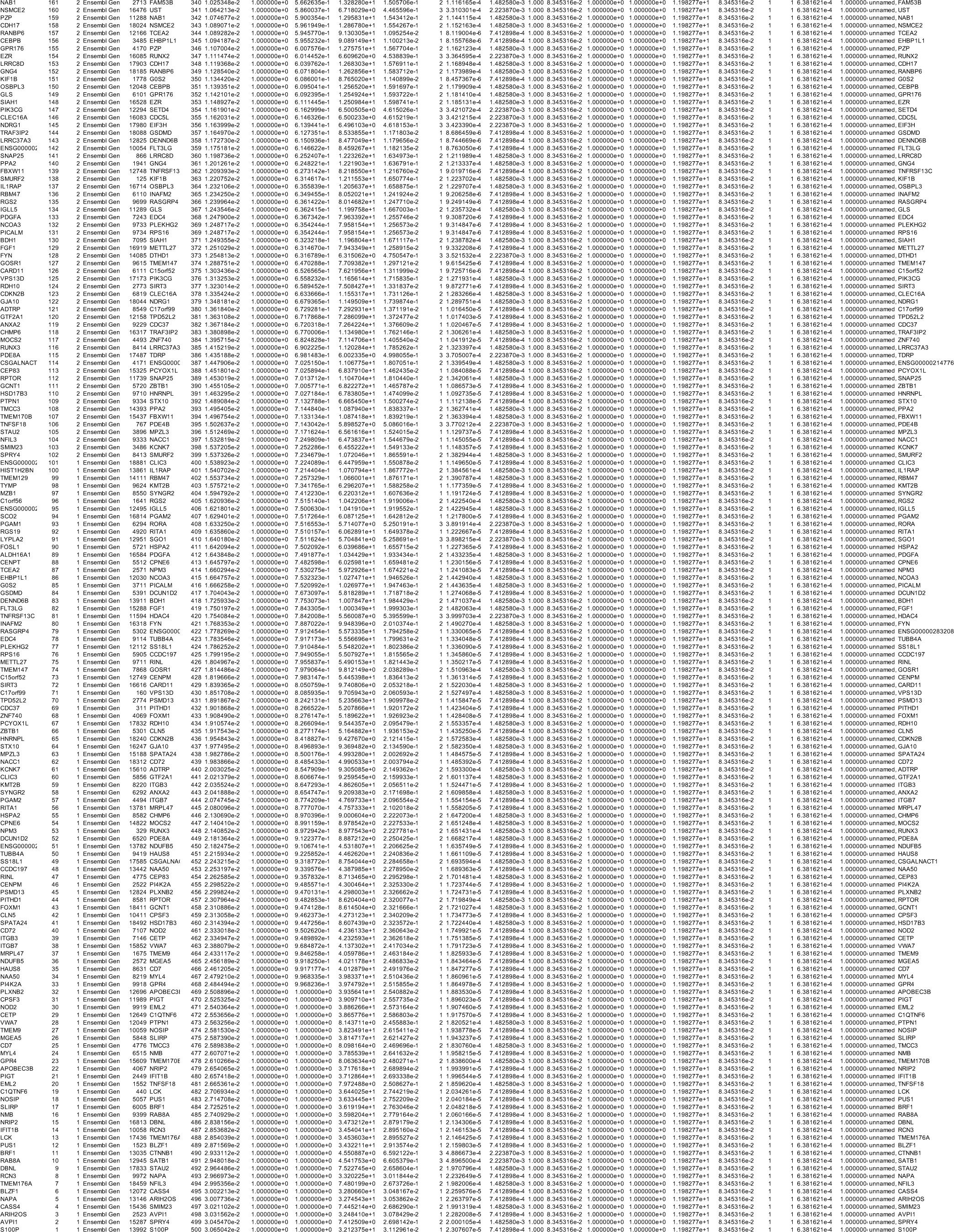
GREAT analysis. Related to Figure 3F

**Table S7.**
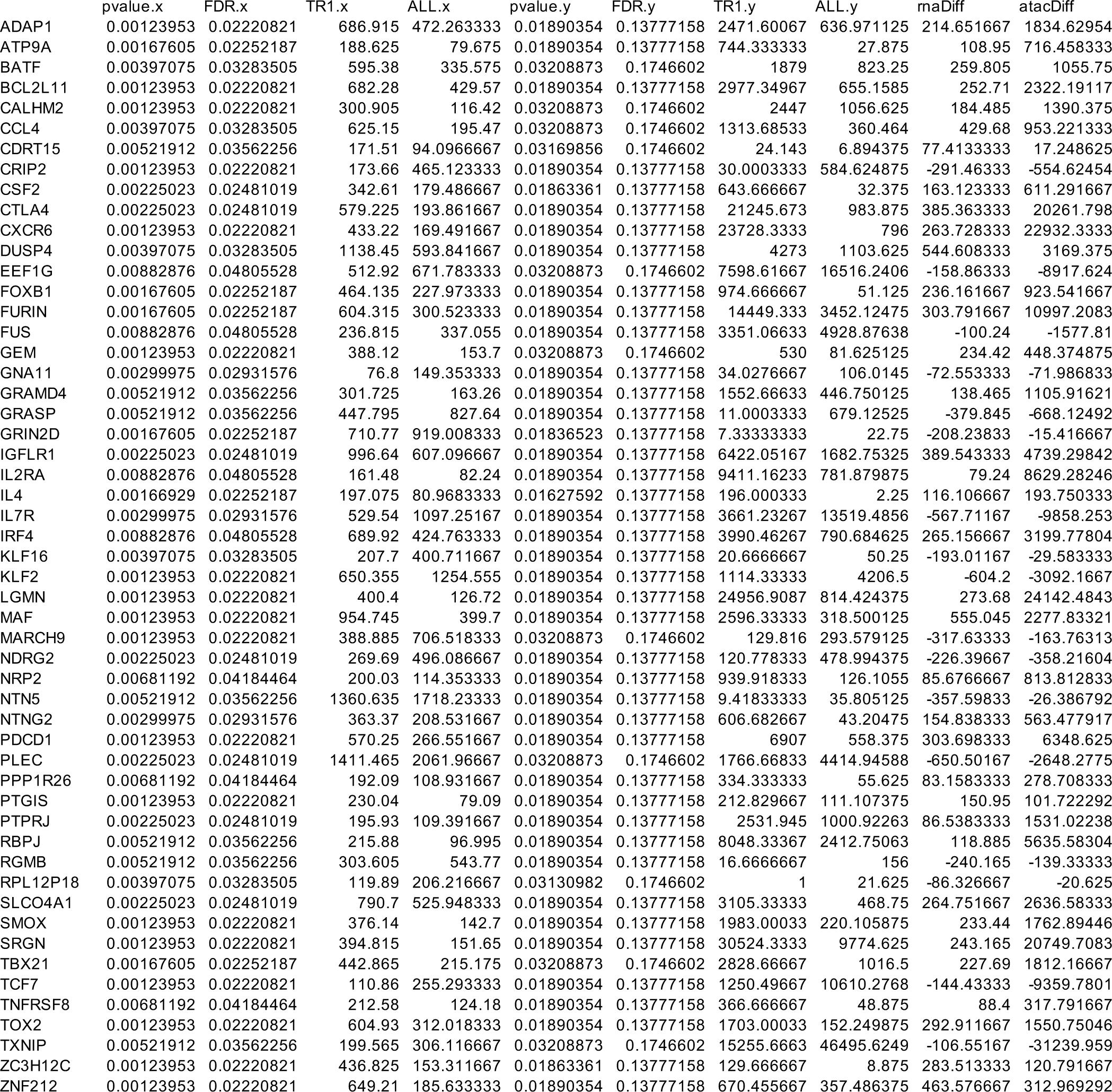
Domains of regulatory chromatin (DORC) analysis. Related to Figure 3G

**Table S8.**
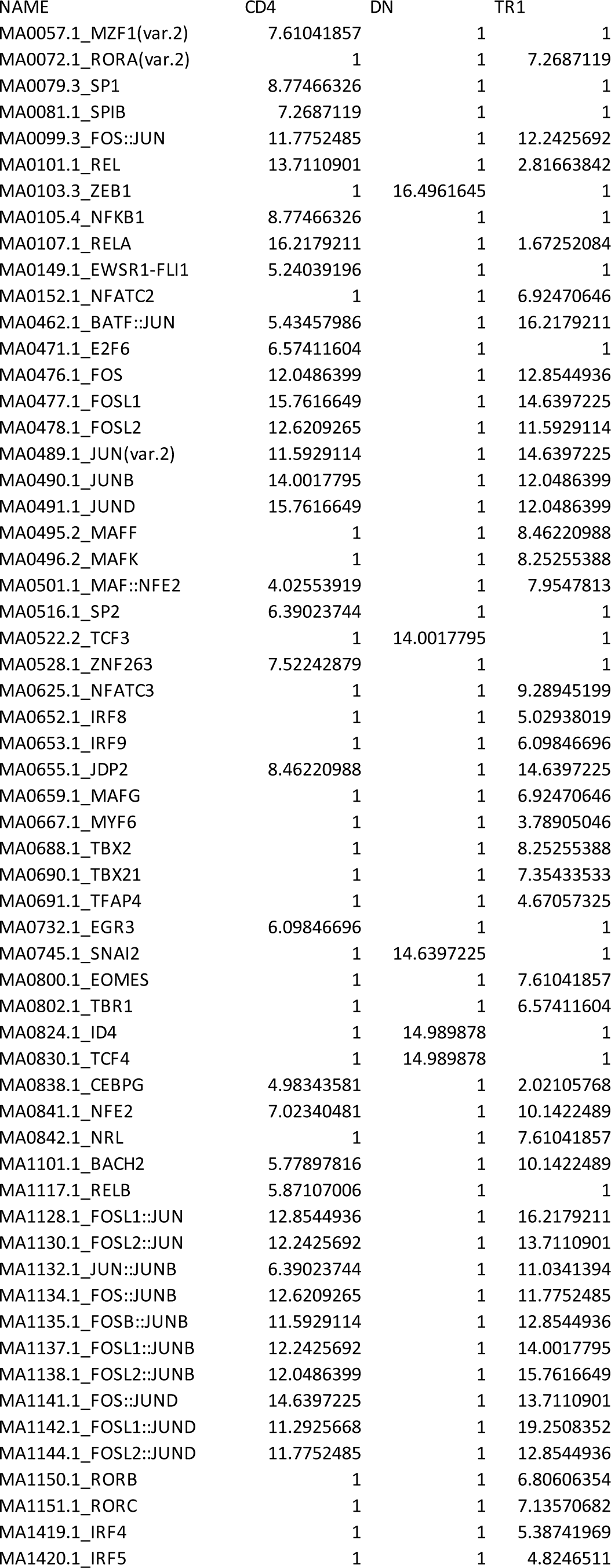
CIBERSORTx signature matrix output. Related to Figure 5

**Table S9.**
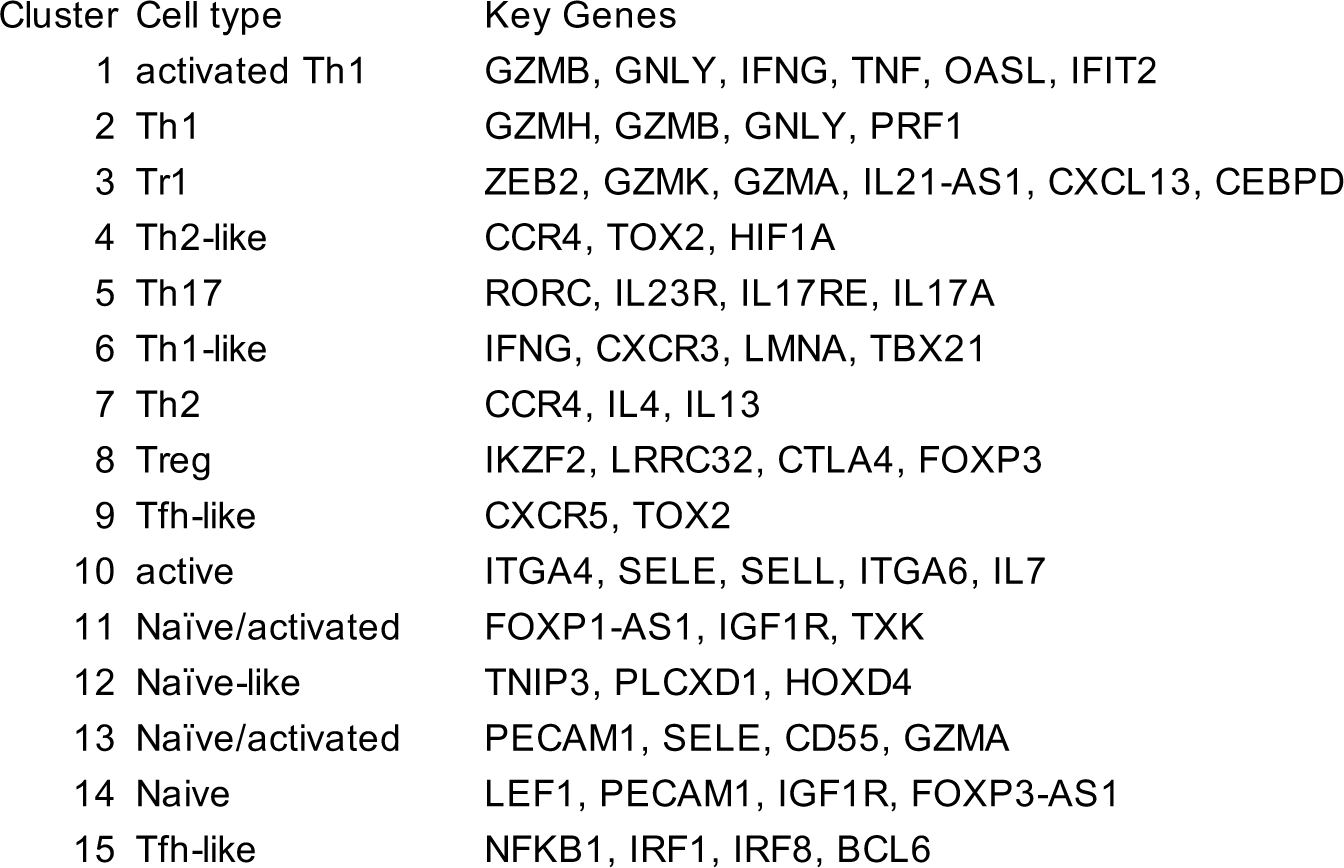
single-cell multiome (ATAC+RNA-seq), cluster annotations. Related to Figure 5

| Cluster | Cell type | Key Genes |
| --- | --- | --- |
| 1 | activated Th1 | GZMB, GNLY, IFNG, TNF, OASL, IFIT2 |
| 2 | Th1 | GZMH, GZMB, GNLY, PRF1 |
| 3 | Tr1 | ZEB2, GZMK, GZMA, IL21-AS1, CXCL13, CEBPD |
| 4 | Th2-like | CCR4, TOX2, HIF1A |
| 5 | Th17 | RORC, IL23R, IL17RE, IL17A |
| 6 | Th1-like | IFNG, CXCR3, LMNA, TBX21 |
| 7 | Th2 | CCR4, IL4, IL13 |
| 8 | Treg | IKZF2, LRRC32, CTLA4, FOXP3 |
| 9 | Tfh-like | CXCR5, TOX2 |
| 10 | active | ITGA4, SELE, SELL, ITGA6, IL7 |
| 11 | Naïve/activated | FOXP1-AS1, IGF1R, TXK |
| 12 | Naïve-like | TNIP3, PLCXD1, HOXD4 |
| 13 | Naïve/activated | PECAM1, SELE, CD55, GZMA |
| 14 | Naïve | LEF1, PECAM1, IGF1R, FOXP3-AS1 |
| 15 | Tfh-like | NFKB1, IRF1, IRF8, BCL6 |

**Table S10.**
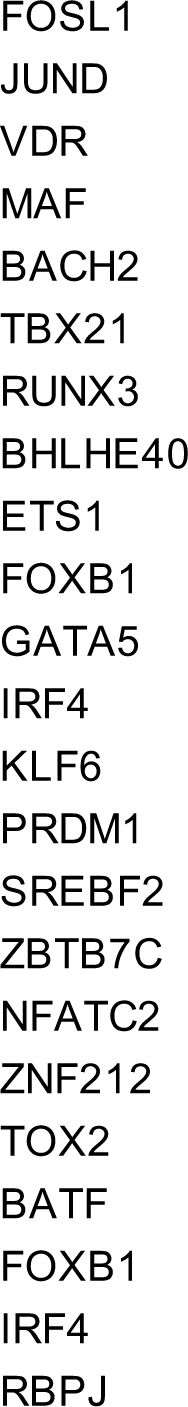
Tr1 TF signature (GSEA gene set matrix) Related to Figure 5

**Table S11.** GSEA analysis of peripheral blood CD4+ T cell single-cell multiome dataset. Related to Figure 5

| Cluster | NES | q-value | NES x -log10(q-value) |
| --- | --- | --- | --- |
| C1 | 1.5415 | 0.0847 | 1.65 |
| C2 | 1.3562 | 0.1834 | 1.00 |
| **C3** | **1.6082** | **0.0402** | **2.24** |
| C4 | 0.8004 | 0.6764 | 0.14 |
| C5 | 1.2763 | 0.23702 | 0.80 |
| C6 | 1.2403 | 0.2815 | 0.68 |
| C7 | 1.5189 | 0.0995 | 1.52 |
| C8 | -0.9922 | 0.42806 | -0.37 |
| C9 | 0.6846 | 0.7784 | 0.07 |
| C10 | -1.2752 | 0.2131 | -0.86 |
| C11 | -1.13302 | 0.3177 | -0.56 |
| C12 | -0.9051 | 0.5356 | -0.25 |
| C13 | -1.3509 | 0.1731 | -1.03 |
| C14 | -1.6437 | 0.0512 | -2.12 |
| C15 | -1.0239 | 0.4198 | -0.39 |

**Table S12.** Marker genes, kidney cancer CD4+ T cell clusters (differential accesibility, FDR <= 0.05 & Log2FC >= 0.5) Related to Figure 6

| Cluster | Mostly in | Representative genes | Annotation |
| --- | --- | --- | --- |
| C1 | PBMC | IKZF2, LRRC32, CTLA4, TIGIT | FOXP3 Tregs |
| C2 | TUMOR | IKZF2, CCR8, IL1R2, CTLA4, BATF | FOXP3 Tregs |
| C3 | PBMC & TUMOR | GZMB, PRF1, GZMH, GNLY, IFNG, CX3CR1 | Th1 |
| C4 | PBMC | IFNL1, OASL, IFNB1, IFNW1, IFIT2, RORC, IL23R | IFN-activated/Th17-like |
| C5 | TUMOR | IL2, IL7R, NFKB1, TNFAIP3, RORC, IL17RE | Th17-like effector |
| C6 | TUMOR | BHLHE40, RBPJ, TGFB1, ZEB2, CCR5, DUSP4, GZMK, EOMES, CEBPD | Tr1 |
| C7 | PBMC & TUMOR | CCR6, RORC, RORA, IL23R, IL22 | Th17 |
| C8 | PBMC | IL4, IL13 | Th2 |
| C9 | PBMC | LEF1, SATB1, PECAM1, SELL | Naïve |
| C10 | TUMOR | CCL23, CCL15, CCL18, CCL4L1 | Chemokine-rich, Tfh-like |
| C11 | PBMC | CCR4, IL4 | Th2-like |
| C12 | PBMC | CXCR5, TIGIT, TOX2 | Tfh |

**Table S13.** GSEA analysis of CD4+ T cell single-cell ATAC data from kidney cancer. Related to Figure 6

| Cluster | NES | q-value | NES x -log10(q-value) |
| --- | --- | --- | --- |
| C1 | -1.1518 | 0.3267 | -0.56 |
| C2 | 0.8344 | 0.63002 | 0.17 |
| C3 | 0.8991 | 0.5717 | 0.22 |
| C4 | 0.8707 | 0.5831 | 0.20 |
| C5 | 1.45406 | 0.1022 | 1.44 |
| **C6** | **2.1371** | **0.001** | **6.41** |
| C7 | 0.6019 | 0.9026 | 0.03 |
| C8 | -0.7111 | 0.7602 | -0.08 |
| **C9** | **-1.6271** | **0.0269** | **-2.55** |
| C10 | 0.7979 | 0.71306 | 0.12 |
| C11 | -1.5981 | 0.0526 | -2.04 |
| C12 | -1.4435 | 0.1003 | -1.44 |

**Table S14.**
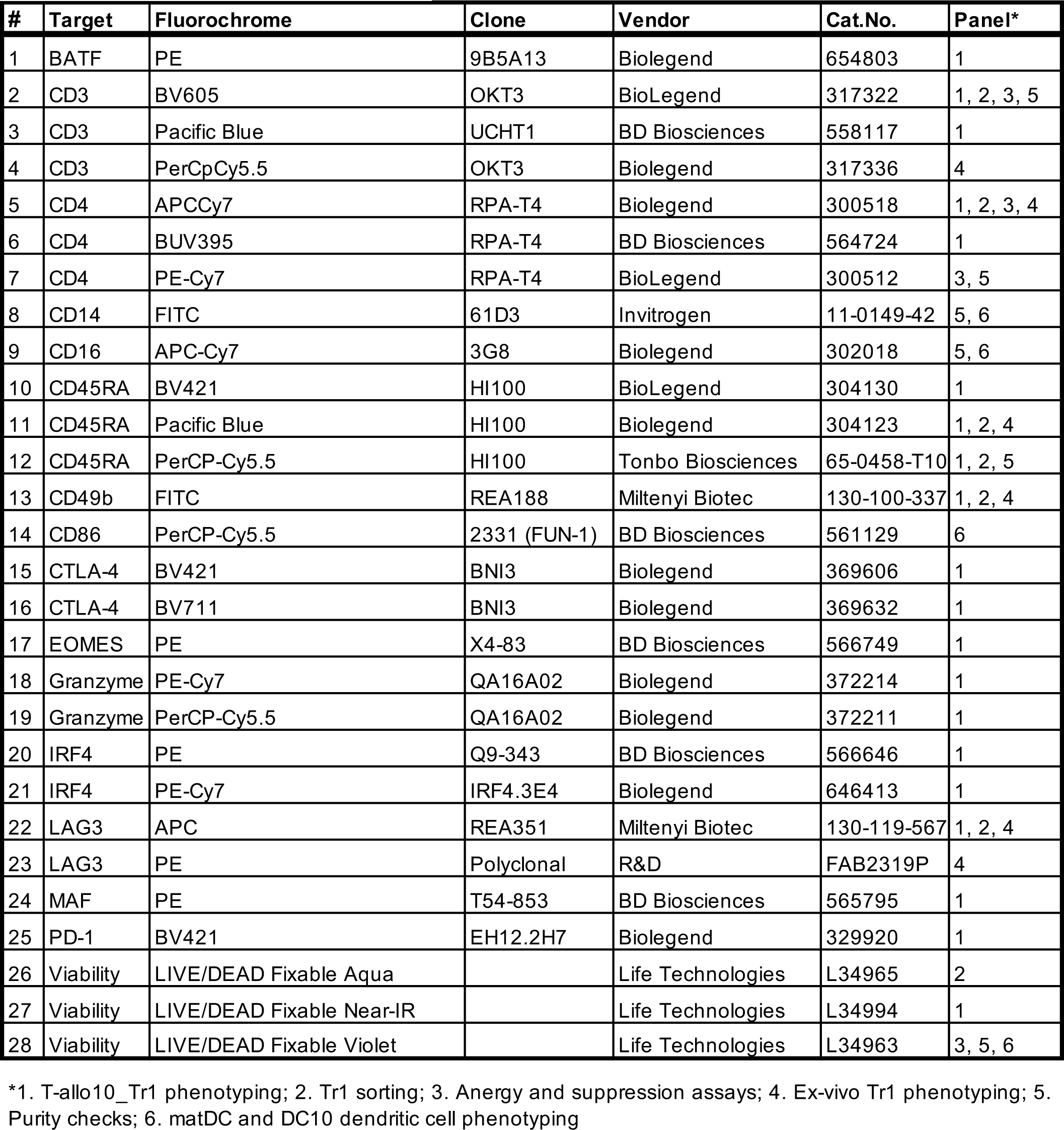
Antibody panels.

**Table S15.**
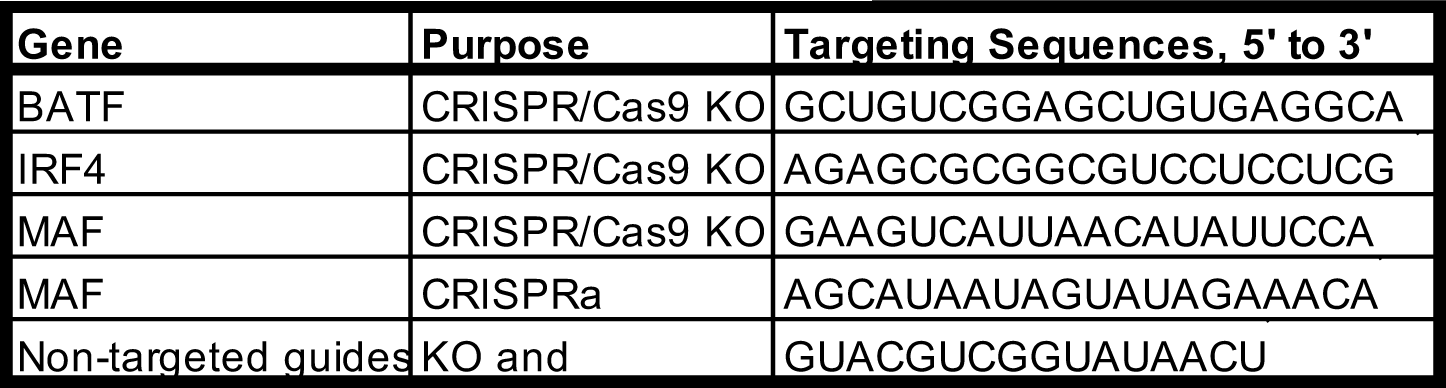
CRISPR guides.

**Table S16.**
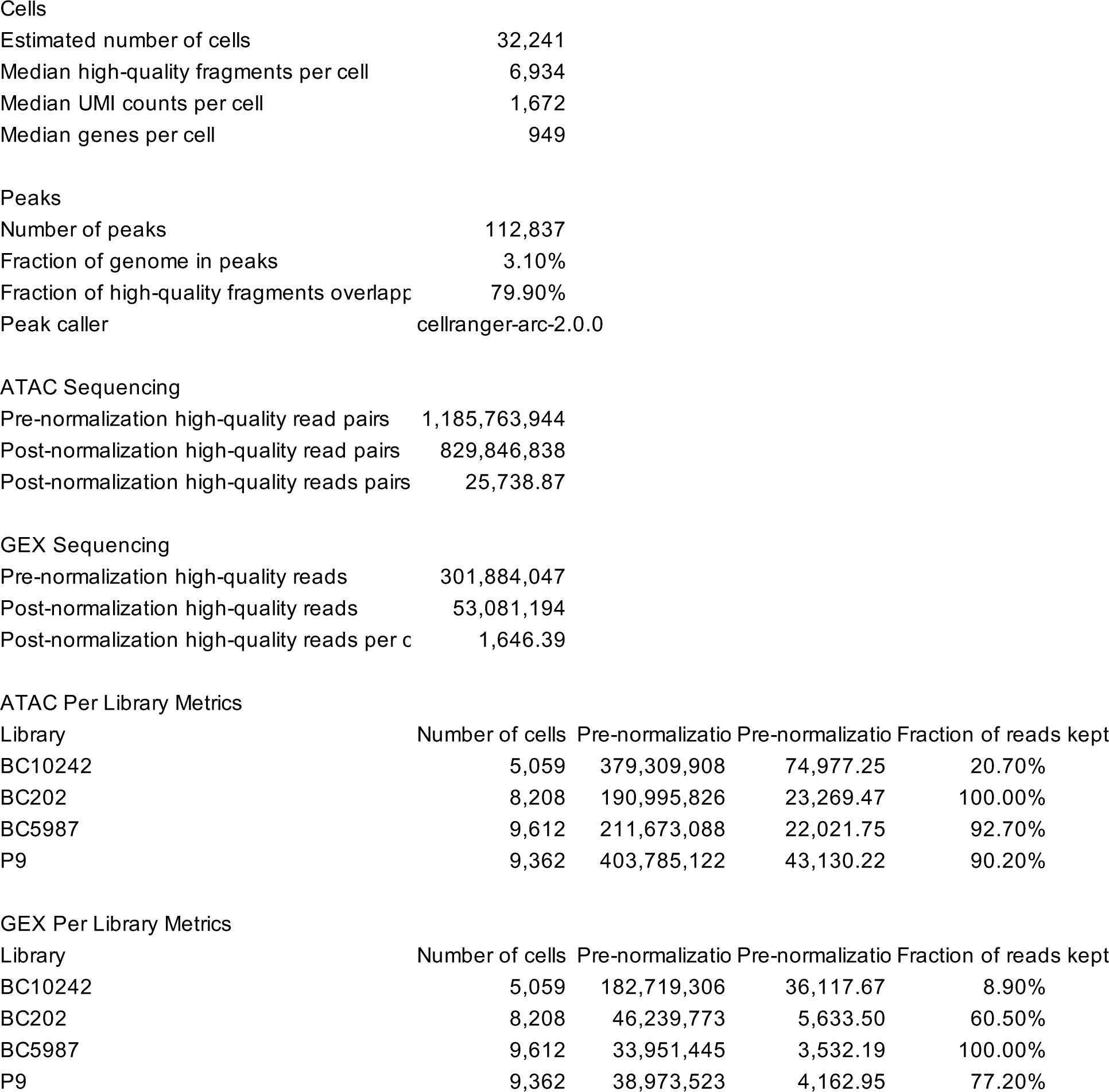
CellRanger scATAC and scRNA-seq ARC output QC.

|  |  |
| --- | --- |
| Estimated number of cells | 32,241 |
| Median high-quality fragments per cell | 6,934 |
| Median UMI counts per cell | 1,672 |
| Median genes per cell | 949 |
| **Peaks** |  |
| Number of peaks | 112,837 |
| Fraction of genome in peaks | 3.10% |
| Fraction of high-quality fragments overlapp | 79.90% |
| Peak caller | cellranger-arc-2.0.0 |
| **ATAC Sequencing** |  |
| Pre-normalization high-quality read pairs | 1,185,763,944 |
| Post-normalization high-quality read pairs | 829,846,838 |
| Post-normalization high-quality reads pairs | 25,738.87 |
| **GEX Sequencing** |  |
| Pre-normalization high-quality reads | 301,884,047 |
| Post-normalization high-quality reads | 53,081,194 |
| Post-normalization high-quality reads per c | 1,646.39 |

**ATAC Per Library Metrics**
| Library | Number of cells | Pre-normalizatio | Pre-normalizatio | Fraction of reads kept |
| --- | --- | --- | --- | --- |
| BC10242 | 5,059 | 379,309,908 | 74,977.25 | 20.70% |
| BC202 | 8,208 | 190,995,826 | 23,269.47 | 100.00% |
| BC5987 | 9,612 | 211,673,088 | 22,021.75 | 92.70% |
| P9 | 9,362 | 403,785,122 | 43,130.22 | 90.20% |

**GEX Per Library Metrics**
| Library | Number of cells | Pre-normalizatio | Pre-normalizatio | Fraction of reads kept |
| --- | --- | --- | --- | --- |
| BC10242 | 5,059 | 182,719,306 | 36,117.67 | 8.90% |
| BC202 | 8,208 | 46,239,773 | 5,633.50 | 60.50% |
| BC5987 | 9,612 | 33,951,445 | 3,532.19 | 100.00% |
| P9 | 9,362 | 38,973,523 | 4,162.95 | 77.20% |

